# Mating status-dependent dopaminergic modulation of auditory sensory neurons in *Drosophila*

**DOI:** 10.1101/2025.05.22.655009

**Authors:** Haruna Yamakoshi, Mihoko Horigome, Shotaro Yamamoto, Shoya Iwanami, Shingo Iwami, Ryoya Tanaka, Yuki Ishikawa, Azusa Kamikouchi

## Abstract

Mating status often modulates responses to courtship sounds in animals. The neural mechanisms underlying this modulation, however, have not been well clarified. Here, we show that dopaminergic signals are involved in modulating the responses of auditory sensory neurons in *Drosophila melanogaster* females depending on their mating status. These neurons abundantly express three types of dopamine receptors, with some having direct synaptic connections with dopaminergic neurons. Of these receptors, suppressing the expression of Dop1R2 reduces sound responses of auditory sensory neurons in unmated but not mated females. Moreover, expression of Dop1R2 in auditory sensory neurons enhances the song response behavior of unmated females, manifested by copulation receptivity when exposed to songs. Our research suggests that dopaminergic modulation via Dop1R2 is involved in mating state-dependent regulation of auditory sensory processing.

## I. Introduction

Mating states often modulate the auditory processing of animals. In green treefrogs (*Hyla cinerea*), neural responses in the auditory midbrain to mating calls are reduced in recently mated females compared to unmated females,^1^ aligning a report that post-reproductive females did not respond to mating calls.^2^ In female toadfish (*Porichthys notatus*), both the sensitivity of auditory hair cells and the behavioral response to male courtship vocalizations increases during reproductive compared to non-reproductive state.^3,4^ Flexible auditory processing should benefit from effectively utilizing limited brain resources by biasing sound representation in a state-dependent manner.^5,6^ However, the mechanisms responsible for mating status-dependent auditory modulation remain poorly understood.

The acoustic communication system of the fruit fly *Drosophila melanogaster* has been used as a model for studying the mechanism of auditory processing (e.g. Baker et al.^7^; Deutsch et al.^8^; Kamikouchi et al.^9^; K. Wang et al.^10^). During mating rituals, *Drosophila* males produce courtship songs by wing vibrations to attract females. Amongst several song types, the pulse song is the primary component for mating success: It carries a species-specific temporal pattern^11^ that selectively enhances female copulation receptivity.^12,13^ After mating, females exhibit decreased copulation receptivity, even in the presence of courtship from conspecific males,^14^ suggesting a reduced response to courtship signals.

Fruit flies detect sound signals with their Johnston’s organs (JOs) located in the second antennal segment.^15^ The JO contains mechanosensory neurons, known as JO neurons, which project their axons to the antennal mechanosensory and motor center (AMMC) located at the ventrolateral side of the brain.^16^ Sound information from the courtship song is primarily transmitted to the AMMC and then relayed to higher-order centers for further processing. Higher-order neurons in the brain, such as AMMC-B1, pC1, pC2l, and vpoEN neurons, are involved in this processing, ultimately modulating the female’s mating decision.^8–10,17,18^ While the neurons comprising the auditory pathway that processes song information are well characterized, it remains unclear whether the properties of this pathway undergo post-mating modulation. The first step in evaluating this is to investigate whether JO neurons, the initial component of the auditory pathway, receive mating status-dependent modulation.

JO neurons, comprising approximately 480 mechanosensory neurons in *D. melanogaster*, are subdivided into five groups, i.e., JO-A to E, each exhibiting distinct response properties and spatially segregated axonal projections in the brain.^9,16^ Among them, JO-A and JO-B (JO-AB hereafter) neurons preferentially respond to antennal vibrations, serving as the major auditory sensory neurons.^9,19,20^ We previously identified abundant postsynaptic sites in the axons of JO-AB neurons within the AMMC.^21^ This finding implied that JO-AB neurons receive synaptic inputs at their axons, which could modulate their response properties.

In this study, we aimed to elucidate cellular mechanisms underlying the mating-dependent auditory modulation in female flies. Using a single-nucleus RNA-seq database, we first examined which neurotransmitter receptors are expressed in JO neurons. Among the identified receptors, we focused on dopamine receptors, as dopamine is used by various animals to modulate sensory information processing in a state-dependent manner. Using T2A-Gal4 knock-in driver lines, we identified three types of dopamine receptors that are abundantly expressed in JO-AB neurons. Combining RNAi-mediated gene knockdown with calcium imaging in unmated and mated females revealed a mating state-dependent effect of Dop1R2 on JO-A responses. Finally, through behavioral assessment, we found that Dop1R2 expression enhances the song response behavior of unmated females. These results demonstrate the potential role of dopamine in linking the mating status to early-stage auditory processing in the female brain.

## II. Results

### Dopamine receptor expression in JO neurons

In fruit flies, major neurotransmitters include acetylcholine, glutamate, GABA, serotonin, dopamine, and octopamine.^22^ To identify which neurotransmitter receptors are expressed in JO neurons, we analyzed the JO neuron dataset derived from published single-nucleus RNA-seq data.^23^ This analysis revealed that multiple types of receptors for each neurotransmitter are expressed, suggesting that JO neurons can receive signals from all six neurotransmitter types (Figure S1A).

Among these neurotransmitters, we focused on dopamine, which plays a role in the state-dependent modulation of sensory information processing across various animals, including mice, flies, and nematodes.^24–26^ Dopamine is also involved in the control of mating drive in males based on their mating status (Zhang et al., 2016^27^, 2021^28^ for flies and mice, respectively), as well as copulation receptivity in females,^29^ suggesting that it may link mating status to neuronal modulation.

The *Drosophila* genome encodes four dopamine receptors: *Dop1R1*,^30^ *Dop1R2*,^31,32^ *Dop2R*^33^ and *DopEcR*.^34^ Analysis of the JO neuron dataset identified the expression of all four types of dopamine receptors, with *DopEcR* being the most abundantly expressed (Figure 1A, Table 1). To validate these analyses, we used T2A-Gal4 knock-in driver lines that mimic dopamine receptor expressions.^35,36^ In JO, *Dop1R1-Gal4* and *DopEcR-Gal4* labeled most cell bodies of JO neurons (about 85.7% and 94.1 % respectively, N=4 and 2 for each driver line; Figures 1B, 1E, S2I, S2L; Table 2; see Figures S2D, S2H for another driver line) while *Dop1R2-Gal4* and *Dop2R-Gal4* labeled fewer (about 22.6% and 15.1 % respectively, N= 4 and 3 for each driver line; Figures 1C, 1D, S2A-C, S2E-G, S2J-K; Table 2).

**Figure 1:**
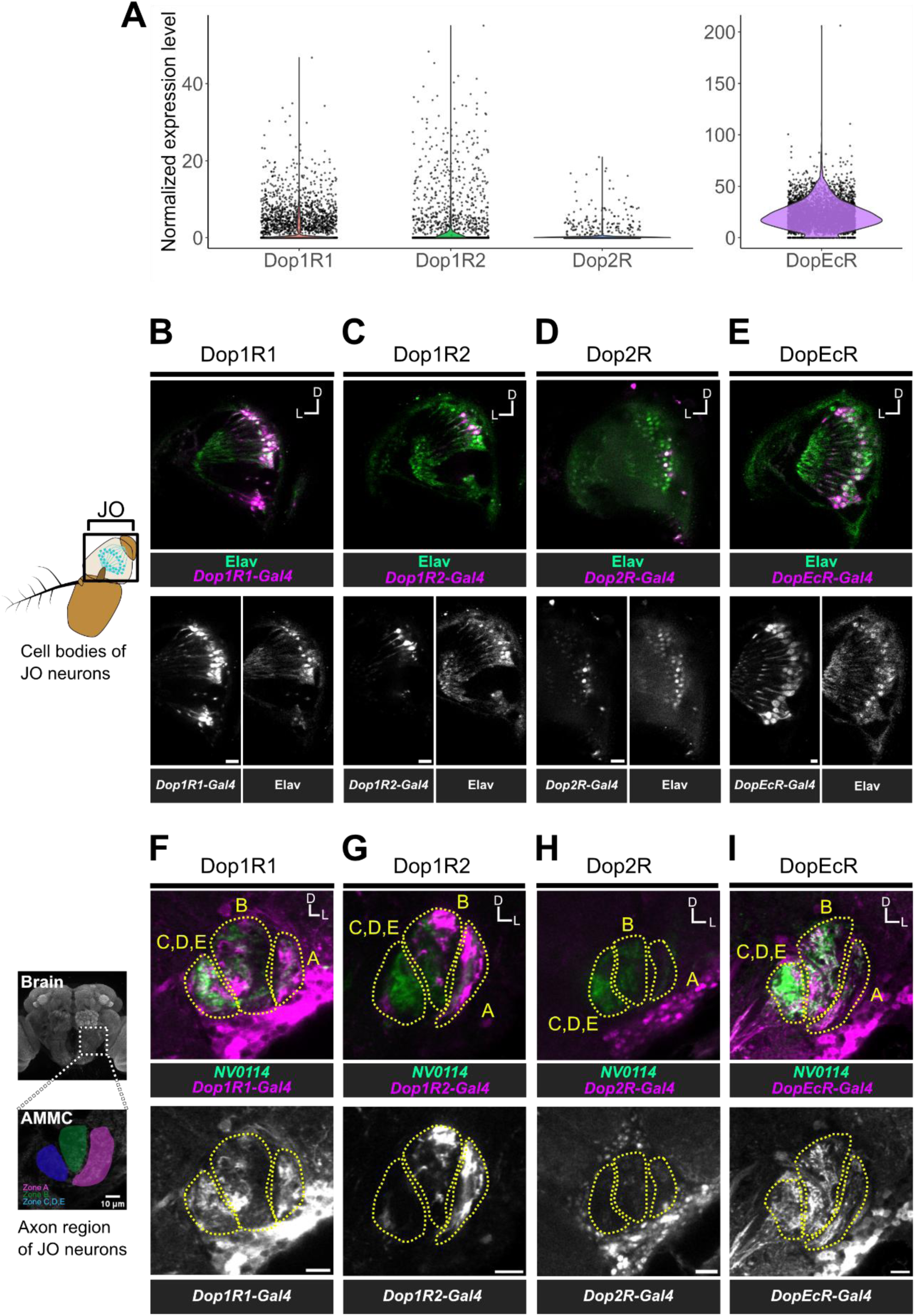
Dopamine receptor expression patterns in JO neurons (A) Expression levels of dopamine receptors in JO neurons. Dots represent the normalized expression level of each gene in single JO neurons obtained from the single-nucleus RNA-seq data.^23^ (B-I) Expression patterns of dopamine receptors in JO (B-E) and AMMC in the brain (F-I). DsRed marker was expressed by T2A-Gal4 knock-in lines to visualize putative dopamine-receptor-expressing cells (magenta). *Dop1R1-Gal4* (B, F), *Dop1R2-Gal4* (C, G), *Dop2R-Gal4* (D, H), and *DopEcR-Gal4* (E, I) (Kondo et al.^36^) were used. Grayscale images of the top panels are shown in the lower panels. Elav (maker of neuronal cell bodies) immunolabeling was used to label JO neurons (green) within the second antennal segment (black square in the left diagram) (B-E). The rCD2::GFP marker expression was driven by the *NV0114* strain, which labels most JO neurons (green) (F-I). JO neurons in each subgroup innervate into five subregions of the AMMC in the brain (yellow dotted lines and the left diagram in F-I). D, dorsal; L, lateral (the same in the following figures). JO, Johnston’s organ; AMMC, antennal mechanosensory and motor center. Scale bar = 10 µm. Genotypes of flies are listed in Table S1 (the same in the following figures).

**Table 1:** Dopamine receptor-positive JO neuron ratio (From single nucleus RNA-seq data)

| Receptor | Proportion of JO neurons |
| --- | --- |
| Dop1R1 | 29.8 % |
| Dop1R2 | 14.4 % |
| Dop2R | 4.11 % |
| DopEcR | 95.2 % |

**Table 2:** Dopamine receptor-positive JO neuron ratio (From immunohistochemistry) Two *Gal4* lines (Deng et al.^35^, D-strain; Kondo et al.^36^, K-strain) were used

| Receptor | AMMC zones | Proportion of JO neurons | Strain | Sample size |
| --- | --- | --- | --- | --- |
| Dop1R1 | A, B, C, D, E | 85.7% | K-strain | 4 |
| Dop1R2 | A, B | 22.6% (Dop1R2-RA) | K-strain | 4 |
|  |  | 15.9% (Dop1R2-RA) | D-strain | 4 |
|  |  | 26.4% (Dop1R2-RC) | D-strain | 3 |
| Dop2R | No expression | 15.1% | K-strain | 3 |
|  | A | 25.3% | D-strain | 3 |
| DopEcR | A, B, C, D, E | 87.6% | D-strain | 2 |
|  |  | 94.1% | K-strain | 2 |

These patterns, together with the RNA-seq database analysis, strongly suggest that all four types of dopamine receptors are expressed in JO neurons but with distinct expression patterns (Table 2): Most JO neurons presumably express *DopEcR* and, less abundantly, *Dop1R2* and *Dop2R*. Although there was a difference in the ratio of *Dop1R1-* positive cells between the RNA-seq data and T2A-Gal4 knock-in driver line (Figures 1A, 1B; Tables 1, 2), both analyses indicate that many JO neurons express *Dop1R1*.

Next, we estimated which subgroups of JO neurons expressed these receptors. JO neurons in each subgroup project their axons to a distinct zone in the AMMC, specifically zones A to E, which correspond to the JO neuron subgroups A to E, respectively (but see Hampel et al.^37^ for another subgroup JO-F; Kamikouchi et al.^16^). This anatomical segregation of axons facilitates the estimation of labeled JO neuron subgroups by examining the labeling patterns of each driver line in the AMMC zones. *Dop1R1-Gal4* and *DopEcR-Gal4* signals were detected broadly across AMMC zones A to E (Figures 1F, 1I, S3D, S3H, S3I, S3L), suggesting that all JO neuron subgroups express these receptors. In contrast, *Dop1R2-Gal4* signals were detected selectively in zones A and B, to which JO-A and JO-B neuron axons project, respectively (Figures 1G, S3A-B, S3E-F, S3J). This observation suggests that Dop1R2 is expressed predominantly in JO-AB neurons. *Dop2R-Gal4* signals were rarely detected (Figures 1H, S3C, S3G, S3K), likely due to low expression levels. Given previous reports suggesting a lack of efferent innervation to JO, which houses the cell bodies of JO neurons,^38^ neurotransmission via these dopamine receptors presumably occurs at the axons of JO neurons in the brain. Overall, we concluded that *Dop1R1* and *DopEcR* are expressed across all subgroups, while *Dop1R2* is expressed selectively in the auditory subset of JO neurons (i.e., JO-AB).

### Dopaminergic neurons synapse to auditory sensory neurons

Dopamine-receptor expression patterns in JO neurons imply direct synaptic connections from dopaminergic neurons. To test this, we utilized a trans-synaptic labeling method, *trans*-Tango (Figure 2A),^39^ driven by two independent *Gal4* strains: One generated by a promoter-Gal4 insertion and the other using a Gal4 cassette knock-in into the tyrosine hydroxylase (TH) coding sequence,^35,40^ both of which presumably label dopaminergic neurons (Figures S4A-B). Either strain induced *trans*-Tango signals in JO neuron cell bodies sparsely (Figures 2B, S4E), with approximately 70 JO neurons showing positive signals on average (Table 3). Given that the *D. melanogaster* JO contains ∼ 480 JO neurons,^16^ a fraction of JO neurons likely receive direct synaptic input from dopaminergic neurons, a proportion much smaller than the population of neurons expressing dopamine receptors (Figure 1; Tables 1, 2; See also in Discussion). In the brain, *trans*-Tango signals were detected sparsely but broadly, spanning the AMMC zones A to E (Figures 2C, S4C-D, S4F). These signals were stronger than those of their parental background flies (i.e., *TH-Gal4* / *w*^1118^ and *w*^1118^ / *trans*-Tango), validating that the observed *trans*-Tango signals in the AMMC are above the autofluorescence level (Figure S4G). These findings suggest that a small subset of JO neurons, encompassing all subgroups, presumably receive synaptic input from dopaminergic neurons.

**Figure 2:**
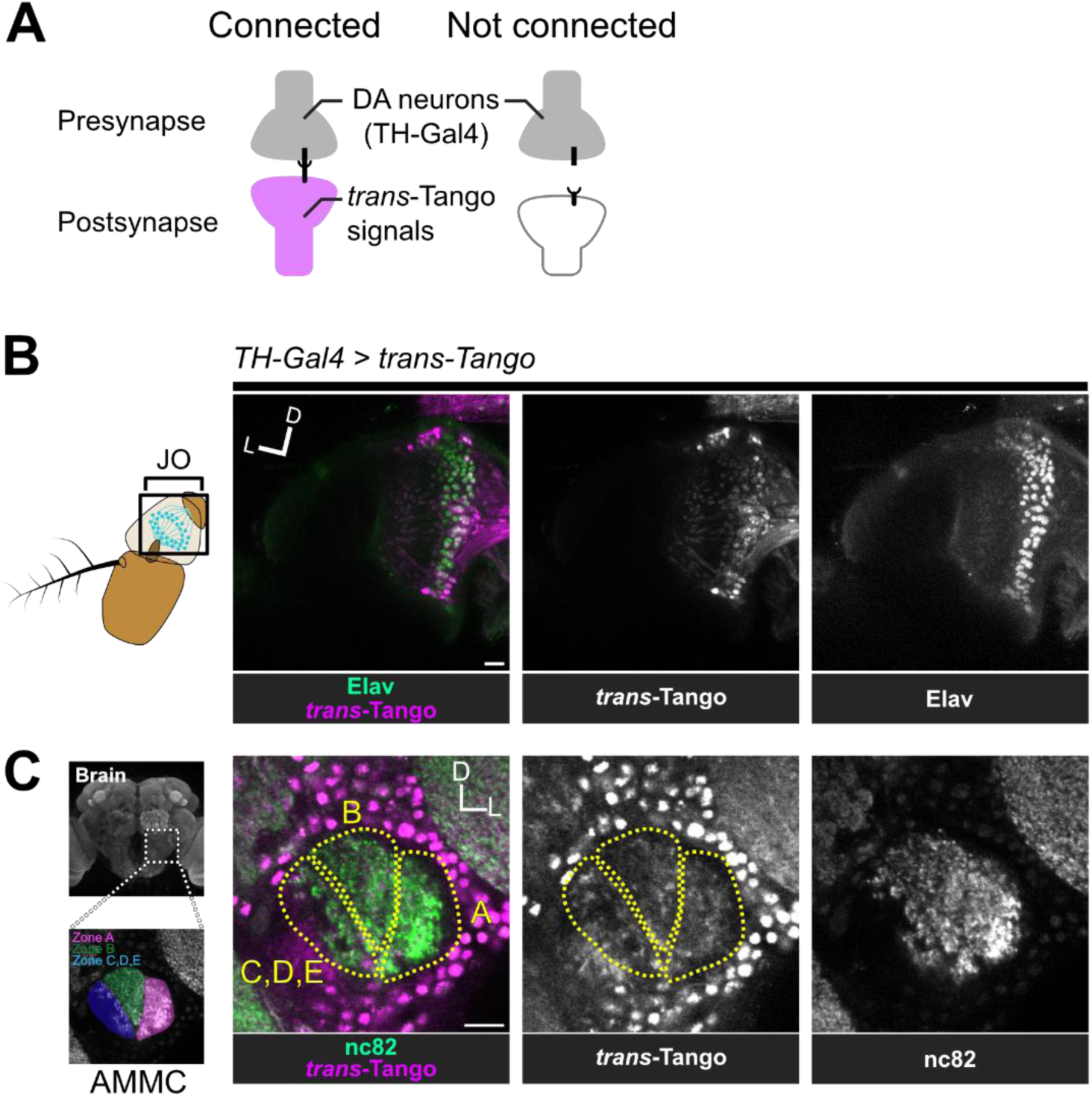
JO neurons have synaptic connections with dopaminergic neurons. (A) *trans*-Tango analysis. *trans*-Tango signals are induced in postsynaptic neurons.^39^ (B-C) *TH-Gal4*^35^ -induced *trans-Tango* signals in JO (B) and AMMC (C). tdTomato was used to visualize *trans*-Tango signals, labeled with anti-DsRed antibodies (magenta). In the JO, cell bodies of JO neurons were labeled with anti-Elav antibodies (green; marker of neuronal cell bodies). In the AMMC, synapses were labeled with nc82 antibodies (green). Scale bar = 10 µm.

**Table 3:**
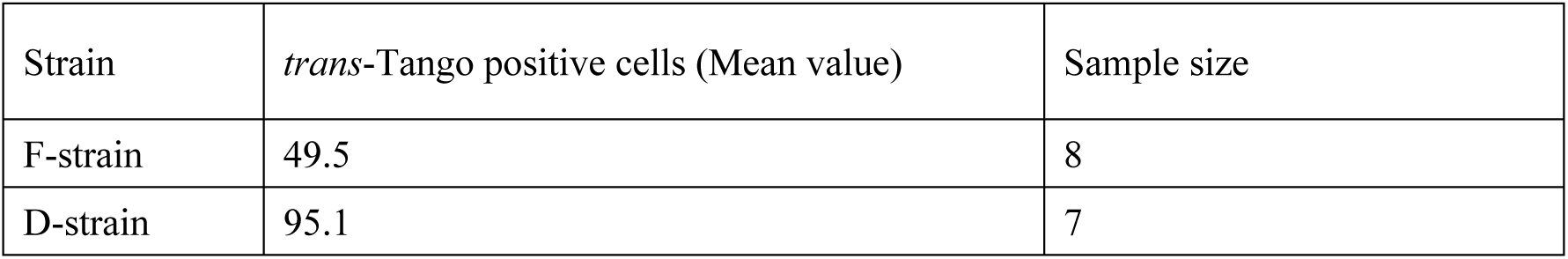
Number of *trans*-Tango positive cells in all JO neurons Two *Gal4* lines (Deng et al.^35^, D-strain; Friggi-Grelin et al.^40^, F-strain) were used

| Strain | *trans*-Tango positive cells (Mean value) | Sample size |
| --- | --- | --- |
| F-strain | 49.5 | 8 |
| D-strain | 95.1 | 7 |

We found that *Drosophila* neuron databases, including Flywire, Flycircuit, and Virtual Fly Brain,^22,41–44^ contain several dopaminergic neurons innervating the AMMC (Tables 4, 5), further supporting our conclusion that dopaminergic neurons have synaptic connections to JO-AB neurons in the brain.

**Table 4:**
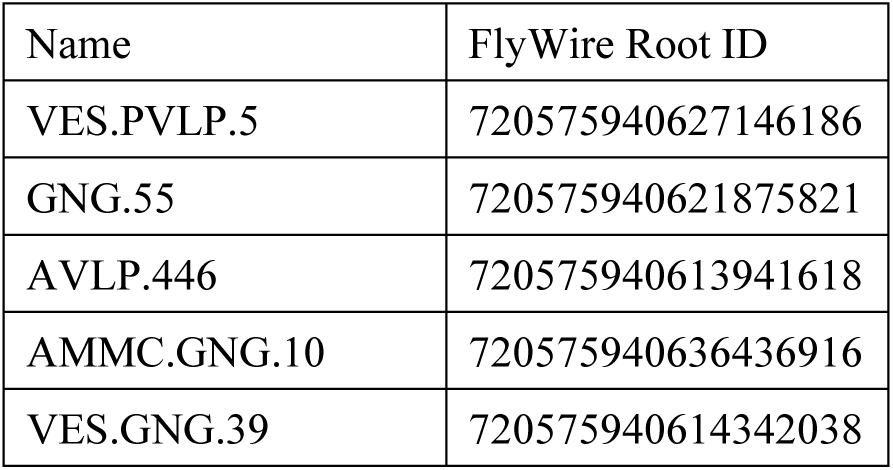
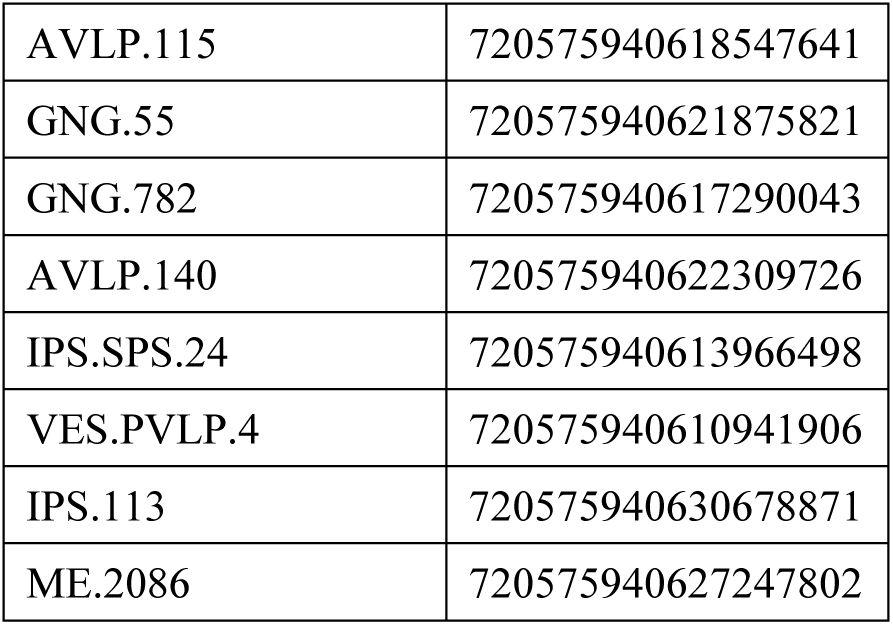
Putative dopaminergic neurons that innervate the AMMC in FlyWire.

| Name | FlyWire Root ID |
| --- | --- |
| VES.PVLP.5 | 720575940627146186 |
| GNG.55 | 720575940621875821 |
| AVLP.446 | 720575940613941618 |
| AMMC.GNG.10 | 720575940636436916 |
| VES.GNG.39 | 720575940614342038 |
| AVLP.115 | 720575940618547641 |
| GNG.55 | 720575940621875821 |
| GNG.782 | 720575940617290043 |
| AVLP.140 | 720575940622309726 |
| IPS.SPS.24 | 720575940613966498 |
| VES.PVLP.4 | 720575940610941906 |
| IPS.113 | 720575940630678871 |
| ME.2086 | 720575940627247802 |

**Table 5:** Putative dopaminergic neurons that innervate zone A or B of the AMMC in FlyCircuit and Virtual Fly Brain.

| FlyCircuit ID | Virtual Fly Brain ID |
| --- | --- |
| TH-F-000025 | VFB_00013621 |
| TH-F-400015 | VFB_00010484 |
| TH-F-200114 | VFB_00013689 |
| TH-F-100058 | VFB_00013576 |
| TH-F-100005 | VFB_00013322 |
| TH-F-000020 | VFB_00013597 |
| TH-F-500006 | VFB_00013699 |
| TH-F-200125 | VFB_00013772 |
| TH-F-200067 | VFB_00013516 |
| TH-F-200029 | VFB_00013465 |
| TH-F-100051 | VFB_00013561 |
| TH-F-100044 | VFB_00013541 |
| TH-F-100031 | VFB_00010473 |
| TH-F-100024 | VFB_00013427 |
| TH-F-100019 | VFB_00013411 |
| TH-F-000100 | VFB_00013766 |
| TH-F-000097 | VFB_00010495 |
| TH-F-000073 | VFB_00013721 |
| TH-F-000051 | VFB_00010483b |

### Dop1R2 expression enhances sound responses of JO-A neurons in unmated females

Next, we aimed to address whether dopamine receptors expressed in JO-AB neurons are involved in their neuronal responses to sound. The contribution of *Dop1R1*, *Dop1R2*, or *DopEcR* expression in JO neurons to the sound-evoked calcium responses was evaluated in unmated (i.e., mature virgin) females, using the following sound stimuli: a 100-Hz pure tone that activates JO-AB strongly,^45^ a pulse song carrying a mean species-specific inter-pulse interval (IPI) (i.e., 35 ms), and a pulse song with a longer IPI (i.e., 75 ms) (Figure 3A). Calcium responses were monitored in the axons of JO neurons, allowing visualization of the responses of each JO neuron subgroup due to the spatial segregation of their axons (Figure 3B).^46^ Although most JO neurons express *Dop1R1* and *DopEcR*, knockdown of these receptor genes in JO neurons had no significant effect on their calcium responses (Table 6; Figure S5). In contrast, knockdown of *Dop1R2*, expressed predominantly in JO-AB neurons, significantly reduced the calcium responses in the axons of JO-A neurons but not of JO-B neurons (Figures 3C-D; Table 6). These decreased responses were observed for all sound stimuli tested, indicating a general reduction in sound responses (p = 0.0107, 0.0107, and 0.00653 for 100 Hz, 35-ms IPI, and 75-ms IPI, respectively; Welch’s t-tests corrected with Benjamini and Hochberg method; Table 6). To evaluate the knockdown effects, we assessed RNAi effectiveness using quantitative RT-qPCR (Figure S6). Each RNAi construct, when driven by the *actin-Gal4 driver*, reduced the expression of the target gene by approximately 47 to 60 % (median values of five repeats) compared to the corresponding controls. By fitting the calcium responses to a simple three-component model (Figure S7), we found that *Dop1R2* knockdown effect in JO-A neurons can be explained by a reduced calcium increase rate during the stimulus (𝑟) and accelerated calcium decrease rate after the stimulus (𝑑_2_) (Figures 3E-G; Table 7). Dop1R2 expression thus likely accelerates the calcium increase during the stimulus and slows the decay afterward in JO-A neurons, thereby enhancing both the overall calcium response and its duration. Altogether, our findings suggest that dopamine signals via Dop1R2 increase the response amplitude of JO-A neurons to sound in unmated females.

**Figure 3:**
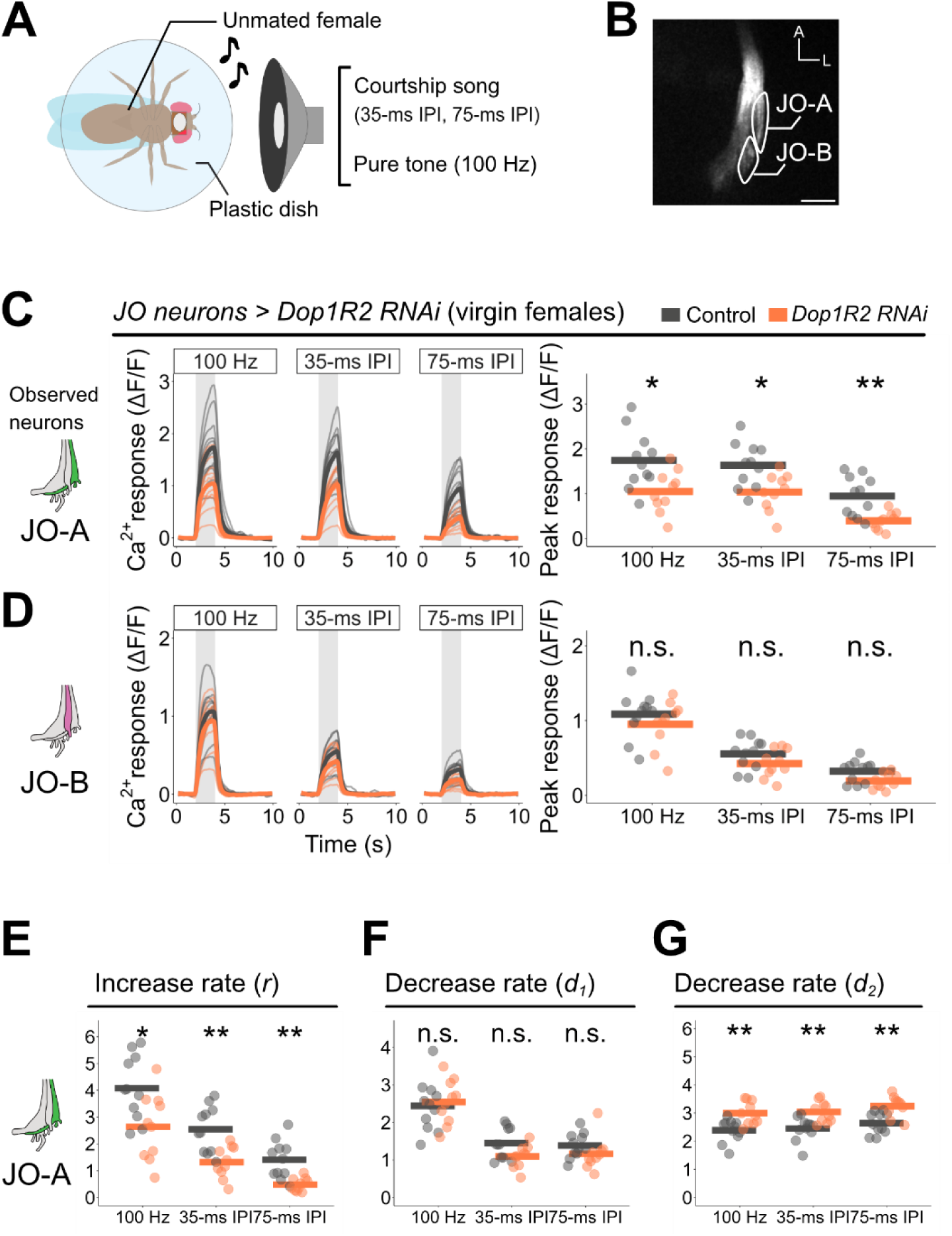
Dop1R2 expression increases JO-A neuron responses in unmated females. (A) Experimental setup of calcium imaging. Unmated females were stabilized ventral side up. Artificial courtship songs (35 ms and 75 ms IPIs) and 100-Hz pure tone were played through a loudspeaker. (B) The region of interest (ROI, outlined in white) for the analysis. Two ROIs were set at the axon bundle of JO-A and JO-B neurons in the AMMC as indicated. A, anterior. Scale bar = 50 µm. (C, D) Calcium responses of JO-A (C) and JO-B (D) in unmated females with *Dop1R2* knockdown in JO neurons. n = 10–11 per genotype. Left, Time traces of raw ΔF/F responses. Thin and bold lines show time traces of the response in each individual and the average of all individuals, respectively. Gray-shaded areas indicate the time window of sound playback. Right, peak calcium responses. Dots and bars show peak responses in each individual and the average of all individuals, respectively (the same in following figures). (E-G) Estimated increase and decrease rates in JO-A neurons. A differential equation that incorporates three factors is fitted to the time series data of calcium responses. 𝑟, estimated increase rate during the stimulus (E); 𝑑_1_, decrease rate during the stimulus (F); 𝑑_2_, decrease rate after the stimulus (G).

**Table 6:**
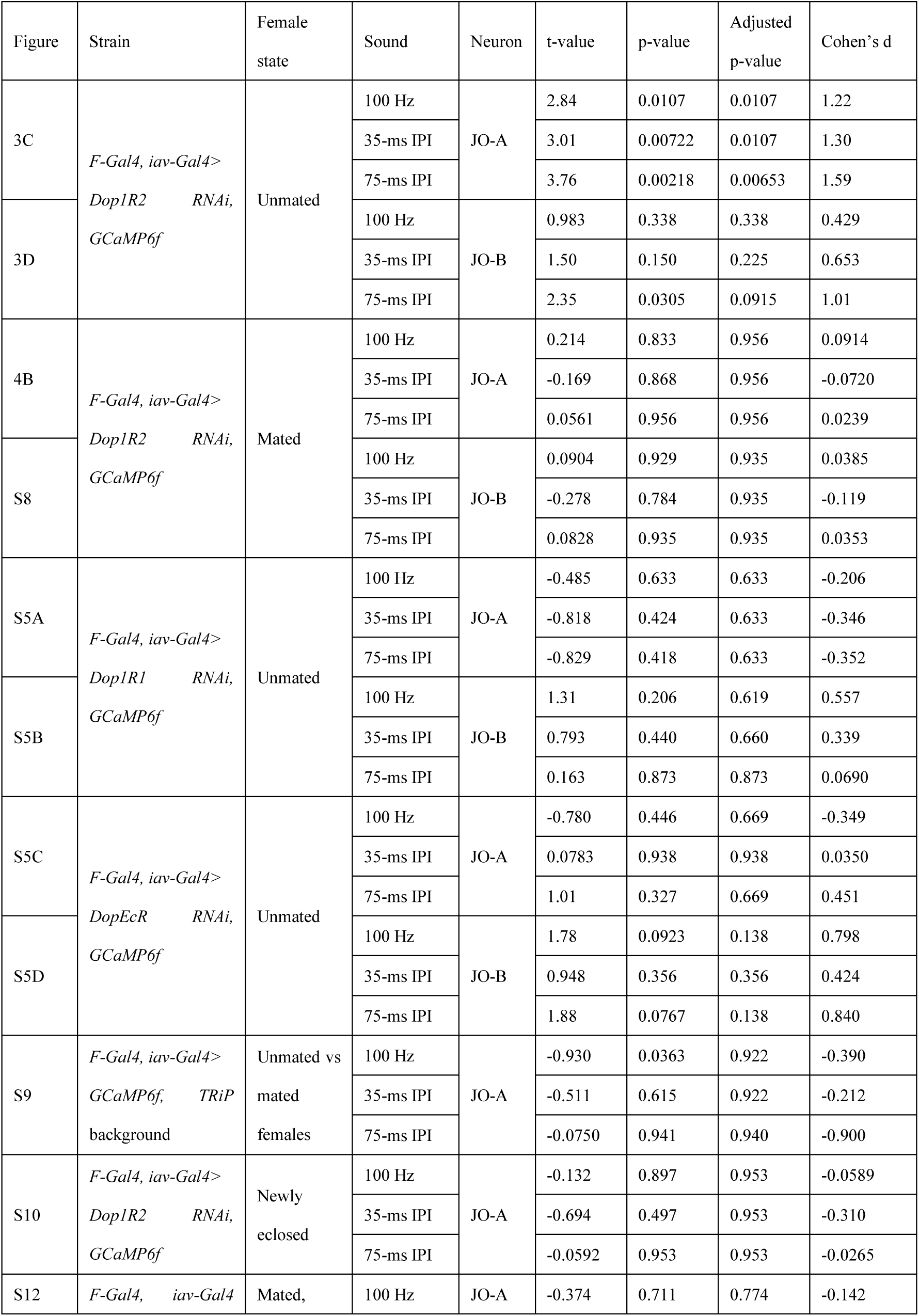
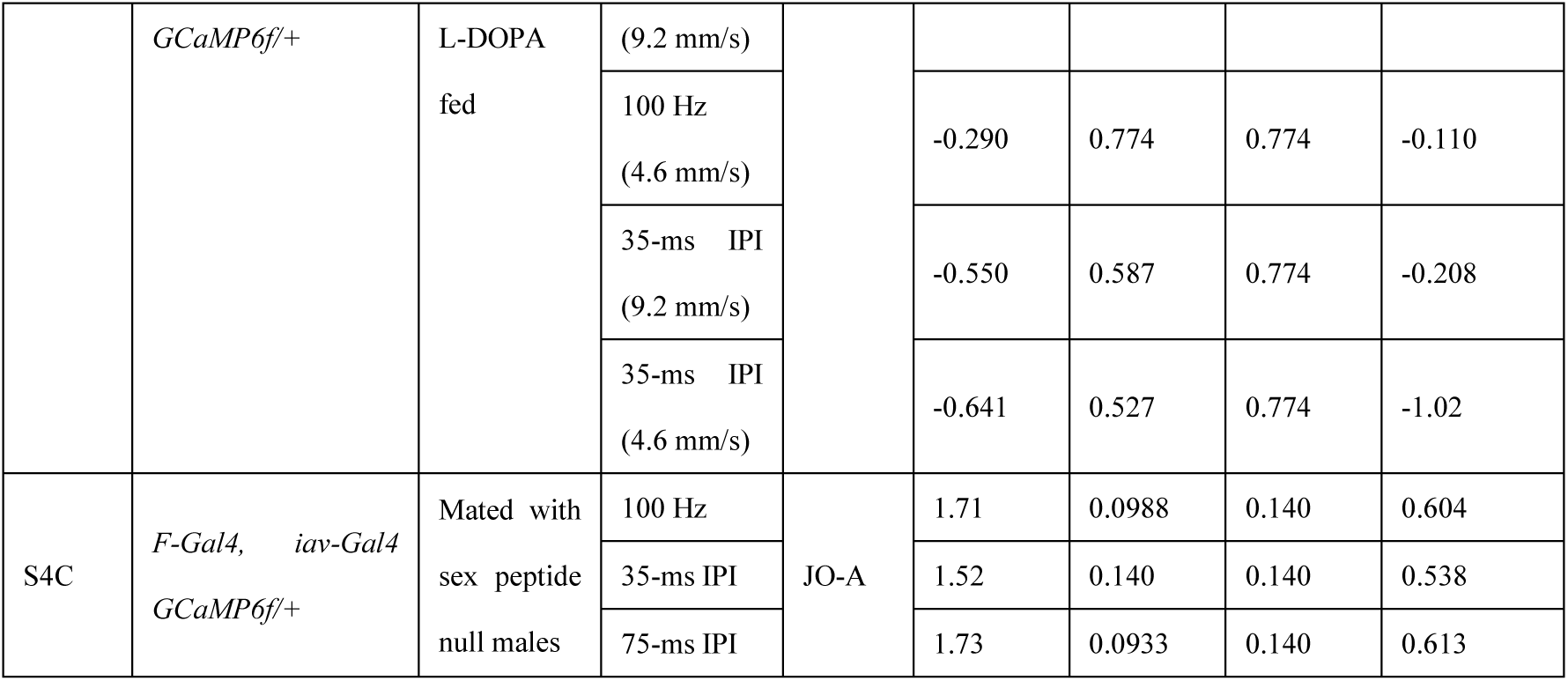
Statistical summary of calcium imaging using Welch’s t-test corrected with Benjamini and Hochberg method.

| Figure | Strain | Female state | Sound | Neuron | t-value | p-value | Adjusted p-value | Cohen's d |
| --- | --- | --- | --- | --- | --- | --- | --- | --- |
| 3C | *F-Gal4, iav-Gal4>*<br>*Dop1R2 RNAi, GCaMP6f* | Unmated | 100 Hz | JO-A | 2.84 | 0.0107 | 0.0107 | 1.22 |
|  |  |  | 35-ms IPI |  | 3.01 | 0.00722 | 0.0107 | 1.30 |
|  |  |  | 75-ms IPI |  | 3.76 | 0.00218 | 0.00653 | 1.59 |
| 3D |  |  | 100 Hz | JO-B | 0.983 | 0.338 | 0.338 | 0.429 |
|  |  |  | 35-ms IPI |  | 1.50 | 0.150 | 0.225 | 0.653 |
|  |  |  | 75-ms IPI |  | 2.35 | 0.0305 | 0.0915 | 1.01 |
| 4B | *F-Gal4, iav-Gal4>*<br>*Dop1R2 RNAi, GCaMP6f* | Mated | 100 Hz | JO-A | 0.214 | 0.833 | 0.956 | 0.0914 |
|  |  |  | 35-ms IPI |  | -0.169 | 0.868 | 0.956 | -0.0720 |
|  |  |  | 75-ms IPI |  | 0.0561 | 0.956 | 0.956 | 0.0239 |
| S8 |  |  | 100 Hz | JO-B | 0.0904 | 0.929 | 0.935 | 0.0385 |
|  |  |  | 35-ms IPI |  | -0.278 | 0.784 | 0.935 | -0.119 |
|  |  |  | 75-ms IPI |  | 0.0828 | 0.935 | 0.935 | 0.0353 |
| S5A | *F-Gal4, iav-Gal4>*<br>*Dop1R1 RNAi, GCaMP6f* | Unmated | 100 Hz | JO-A | -0.485 | 0.633 | 0.633 | -0.206 |
|  |  |  | 35-ms IPI |  | -0.818 | 0.424 | 0.633 | -0.346 |
|  |  |  | 75-ms IPI |  | -0.829 | 0.418 | 0.633 | -0.352 |
| S5B |  |  | 100 Hz | JO-B | 1.31 | 0.206 | 0.619 | 0.557 |
|  |  |  | 35-ms IPI |  | 0.793 | 0.440 | 0.660 | 0.339 |
|  |  |  | 75-ms IPI |  | 0.163 | 0.873 | 0.873 | 0.0690 |
| S5C | *F-Gal4, iav-Gal4>*<br>*DopEcR RNAi, GCaMP6f* | Unmated | 100 Hz | JO-A | -0.780 | 0.446 | 0.669 | -0.349 |
|  |  |  | 35-ms IPI |  | 0.0783 | 0.938 | 0.938 | 0.0350 |
|  |  |  | 75-ms IPI |  | 1.01 | 0.327 | 0.669 | 0.451 |
| S5D |  |  | 100 Hz | JO-B | 1.78 | 0.0923 | 0.138 | 0.798 |
|  |  |  | 35-ms IPI |  | 0.948 | 0.356 | 0.356 | 0.424 |
|  |  |  | 75-ms IPI |  | 1.88 | 0.0767 | 0.138 | 0.840 |
| S9 | *F-Gal4, iav-Gal4>*<br>*GCaMP6f, TRiP*<br>background | Unmated vs mated females | 100 Hz | JO-A | -0.930 | 0.0363 | 0.922 | -0.390 |
|  |  |  | 35-ms IPI |  | -0.511 | 0.615 | 0.922 | -0.212 |
|  |  |  | 75-ms IPI |  | -0.0750 | 0.941 | 0.940 | -0.900 |
| S10 | *F-Gal4, iav-Gal4>*<br>*Dop1R2 RNAi, GCaMP6f* | Newly eclosed | 100 Hz | JO-A | -0.132 | 0.897 | 0.953 | -0.0589 |
|  |  |  | 35-ms IPI |  | -0.694 | 0.497 | 0.953 | -0.310 |
|  |  |  | 75-ms IPI |  | -0.0592 | 0.953 | 0.953 | -0.0265 |
| S12 | *F-Gal4, iav-Gal4* | Mated, | 100 Hz | JO-A | -0.374 | 0.711 | 0.774 | -0.142 |
|  | *GCaMP6f/+* | L-DOPA fed | (9.2 mm/s) |  |  |  |  |  |
|  |  |  | 100 Hz (4.6 mm/s) |  | -0.290 | 0.774 | 0.774 | -0.110 |
|  |  |  | 35-ms IPI (9.2 mm/s) |  | -0.550 | 0.587 | 0.774 | -0.208 |
|  |  |  | 35-ms IPI (4.6 mm/s) |  | -0.641 | 0.527 | 0.774 | -1.02 |
| S4C | *F-Gal4, iav-Gal4*<br>*GCaMP6f/+* | Mated with sex peptide null males | 100 Hz | JO-A | 1.71 | 0.0988 | 0.140 | 0.604 |
|  |  |  | 35-ms IPI |  | 1.52 | 0.140 | 0.140 | 0.538 |
|  |  |  | 75-ms IPI |  | 1.73 | 0.0933 | 0.140 | 0.613 |

**Table 7:**
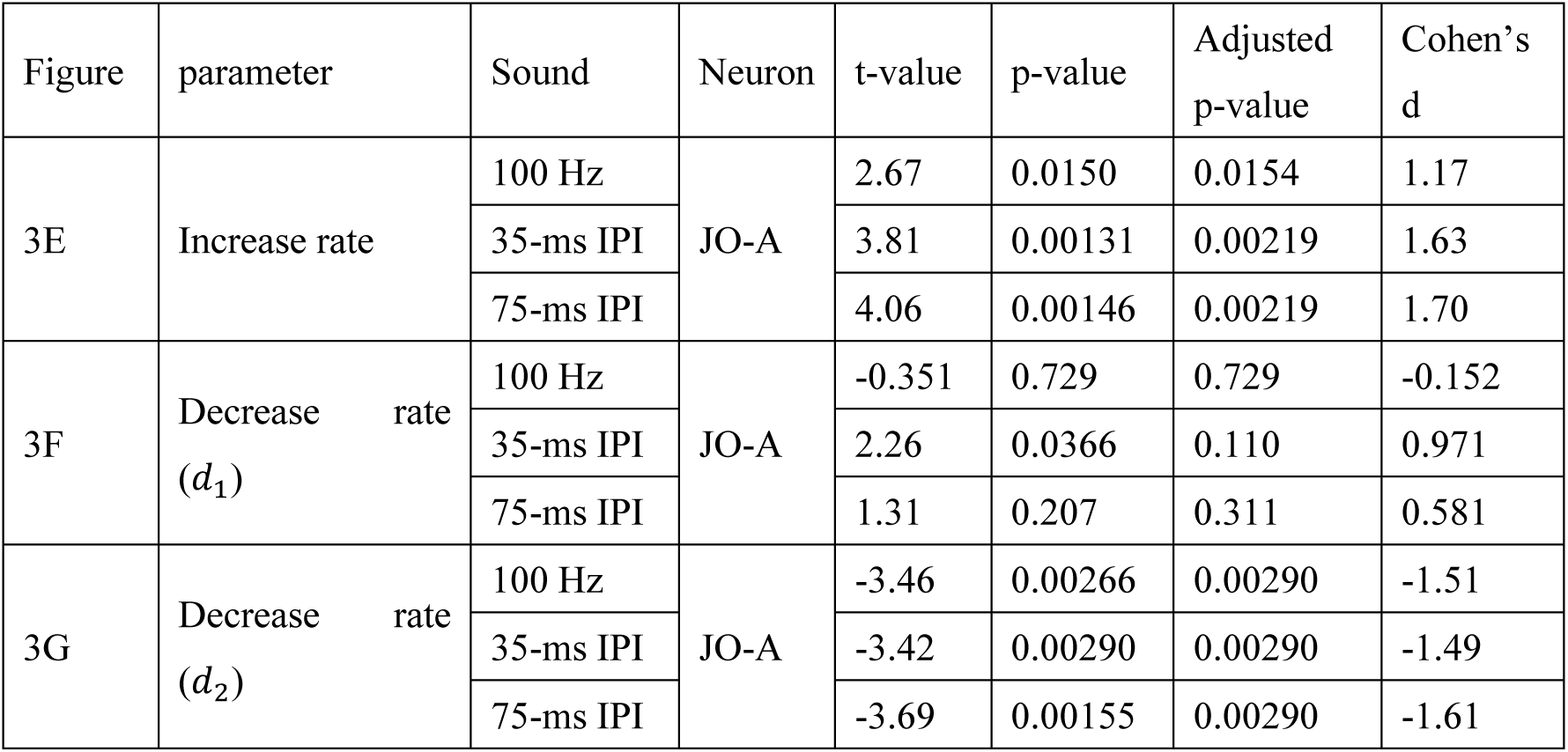
Statistical summary of predicted increase (. 𝑟**) and decrease rates (**𝑑_1_**and** 𝑑_2_)

| Figure | parameter | Sound | Neuron | t-value | p-value | Adjusted p-value | Cohen's d |
| --- | --- | --- | --- | --- | --- | --- | --- |
| 3E | Increase rate | 100 Hz | JO-A | 2.67 | 0.0150 | 0.0154 | 1.17 |
|  |  | 35-ms IPI |  | 3.81 | 0.00131 | 0.00219 | 1.63 |
|  |  | 75-ms IPI |  | 4.06 | 0.00146 | 0.00219 | 1.70 |
| 3F | Decrease rate ($d_1$) | 100 Hz | JO-A | -0.351 | 0.729 | 0.729 | -0.152 |
|  |  | 35-ms IPI |  | 2.26 | 0.0366 | 0.110 | 0.971 |
|  |  | 75-ms IPI |  | 1.31 | 0.207 | 0.311 | 0.581 |
| 3G | Decrease rate ($d_2$) | 100 Hz | JO-A | -3.46 | 0.00266 | 0.00290 | -1.51 |
|  |  | 35-ms IPI |  | -3.42 | 0.00290 | 0.00290 | -1.49 |
|  |  | 75-ms IPI |  | -3.69 | 0.00155 | 0.00290 | -1.61 |

### Dopamine reflects mating status to adjust JO-A responses

To investigate whether Dop1R2-mediated modulation of JO-A neurons depends on the mating status of flies, we examined responses in mated females. Mated females were prepared by housing females with males for 12 to 24 hours prior to calcium imaging (Figure 4A). This procedure resulted in fertilization of over 98% of females, as confirmed by quantifying the population of females that produced larvae after housing with wild-type males. Although JO-A neurons in mated females exhibited robust calcium responses to sounds, they lost susceptibility to *Dop1R2* expression observed in unmated females, as *Dop1R2* knockdown in mated females did not affect the calcium responses of JO-A neurons (p=0.956 for 100 Hz, 35-ms IPI and 75-ms IPI; Welch’s t-test corrected with Benjamini and Hochberg method; Figure 4B; Table 6; see Figure S8 for JO-B responses). A direct comparison of JO-A responses between unmated and mated females revealed no significant difference (p=0.922 for 100 Hz, 35-ms IPI and p=0.940 for 75-ms IPI; Welch’s t-test corrected with Benjamini and Hochberg method; Figure S9; Table 6). This finding suggests that JO-A neurons in unmated females require Dop1R2 expression to reach the response level observed in mated females.

**Figure 4:**
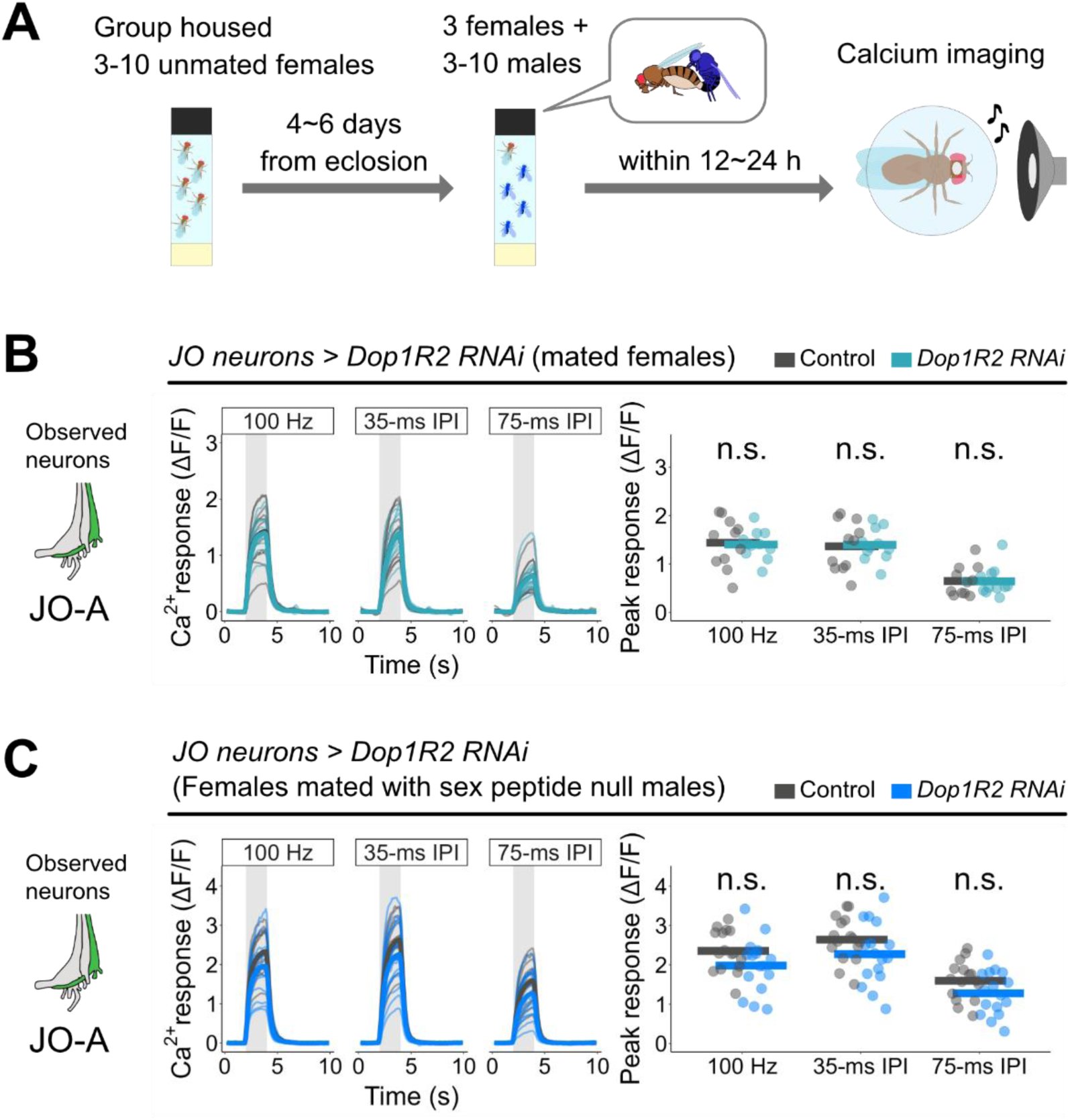
Dop1R2 expression does not affect JO-A responses in mated females. (A) Experimental scheme for calcium imaging in mated females. Females are maintained with males (blue) for 12 to 24 h before being subjected to the calcium imaging. (B) Calcium responses in JO-A axons of mated females. n =11 per genotype. (C) Calcium responses in JO-A axons of pseudo-unmated females (mated with sex peptide null males). n =16 per genotype.

Mated females typically exhibit low copulation receptivity,^14^ suggesting a link between mating receptivity and Dop1R2-mediated modulation. Since newly eclosed females also exhibit low copulation receptivity,^47^ we tested their JO-A responses with *Dop1R2* knockdown. Interestingly, *Dop1R2* knockdown did not significantly affect their calcium responses of JO-A neurons (p=0.953 for 100 Hz, 35-ms IPI and 75-ms IPI; Welch’s t-test corrected with Benjamini and Hochberg method; Figure S10; Table 6). Therefore, together with the results in Figure 3C, these findings suggest that dopamine supports the responses of JO-A neurons via Dop1R2 specifically in females with high mating drive.

The molecular mechanism underlying this status-dependent modulation can be attributed to several steps in dopamine signaling, including receptor amount, dopamine release, and subsequent intracellular signaling pathways. Among these steps, we first examined the amount of receptors located in JO-A axons. We utilized the split-GFP method, where endogenous Dop1R2 is tagged with the C-terminal fragment of the GFP protein (GFP_11_), allowing GFP to be reconstituted when the N-terminal fragment (GFP_1–10_) is expressed in the target neurons.^36^ The reconstituted GFP signals in JO-A axons showed no significant changes between unmated and mated females (p= 0.974, Welch’s t-test; Figure S11; Tables 8, 9), suggesting that receptor abundance in JO-A axons is similar between unmated and mated females. We next examined whether the decrease in dopamine release in mated females underlies this status-dependent modulation. Feeding mated females L-DOPA, a dopamine precursor, did not however alter the calcium responses of JO-A neurons (p=0.774 for 100 Hz, 35-ms IPI and 75-ms IPI, Welch’s t-test corrected with Benjamini and Hochberg method; Figure S12; Table 6). These results propose a model in which the modulation of the Dop1R2 signaling pathway, rather than the expression level of Dop1R2 or the amount of dopamine release, influences the mating-status dependent function of Dop1R2 in JO-A neurons (Figure S13A).

**Table 8:**
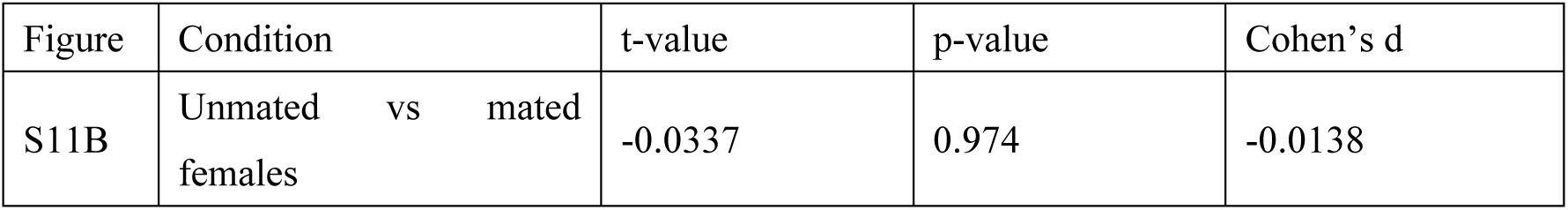
Statistical summary of split-GFP analysis using Welch’s t-test.

**Table 9:** Statistical summary of split-GFP analysis. We also quantified the fluorescence of GFP_1-10_ as GFP_1-10_ itself has fluorescence.^36^ Fluorescence without GFP_11_ was weaker than fluorescence with GFP11

| Figure | Condition | Presence of GFP<sub>11</sub> | Mean | Median | Standard error |
| --- | --- | --- | --- | --- | --- |
| S11B | Unmated | Yes | 526 | 529 | 11.6 |
| S11B | Mated | Yes | 504 | 510 | 22.8 |
| - | Unmated | No (only GFP<sub>1-10</sub>) | 477 | 484 | 16.8 |
| - | Mated | No (only GFP<sub>1-10</sub>) | 452 | 449 | 14.3 |

Finally, we attempted to identify the factors underlying the mating status-dependent, Dop1R2-mediated regulation of JO-A neurons. One promising candidate is sex peptide (SP), which is transferred from males to females during mating and is known to trigger post-mating responses in females.^48–51^ Analysis of the single-nucleus RNA-seq dataset of fruit flies revealed that the receptor for SP is likely expressed in putative JO-AB neurons (Figure S1B). To test whether SP contributes to changes in Dop1R2 function in mated females, we measured calcium responses in pseudo-unmated females that had mated with SP-null males *SP0/ Df(3L)Δ130*.^14^ These pseudo-unmated females exhibited a trend toward reduced JO-A calcium responses when *Dop1R2* was knocked down, although the effect did not reach statistical significance (p=0.139 for 100 Hz, 35-ms IPI and p=0.940 for 75-ms IPI; Welch’s t-test corrected with Benjamini and Hochberg method; Figure 4C; Table 6). These results suggest that SP may have a partial, albeit minor, role in the loss of Dop1R2 dependency in JO-A responses following mating (Figure 4B).

### *Dop1R2* expression in JO-AB neurons affects song response behavior

Dopaminergic modulation of JO neuron responses in unmated females might affect their behavioral responses to sounds. We examined whether dopamine receptors expressed in JO neurons influence the song response behavior of unmated females, which can be measured as copulation receptivity to mute males under artificial song exposure (Figure 5A).^18^ A *F-Gal4* driver line that selectively labels most JO neurons was used for knocking down dopamine receptor expression, although it also labels a few neurons projecting from the legs to the ventral nerve cord (VNC) (Figures 5B, S14A).^9,52^

**Figure 5:**
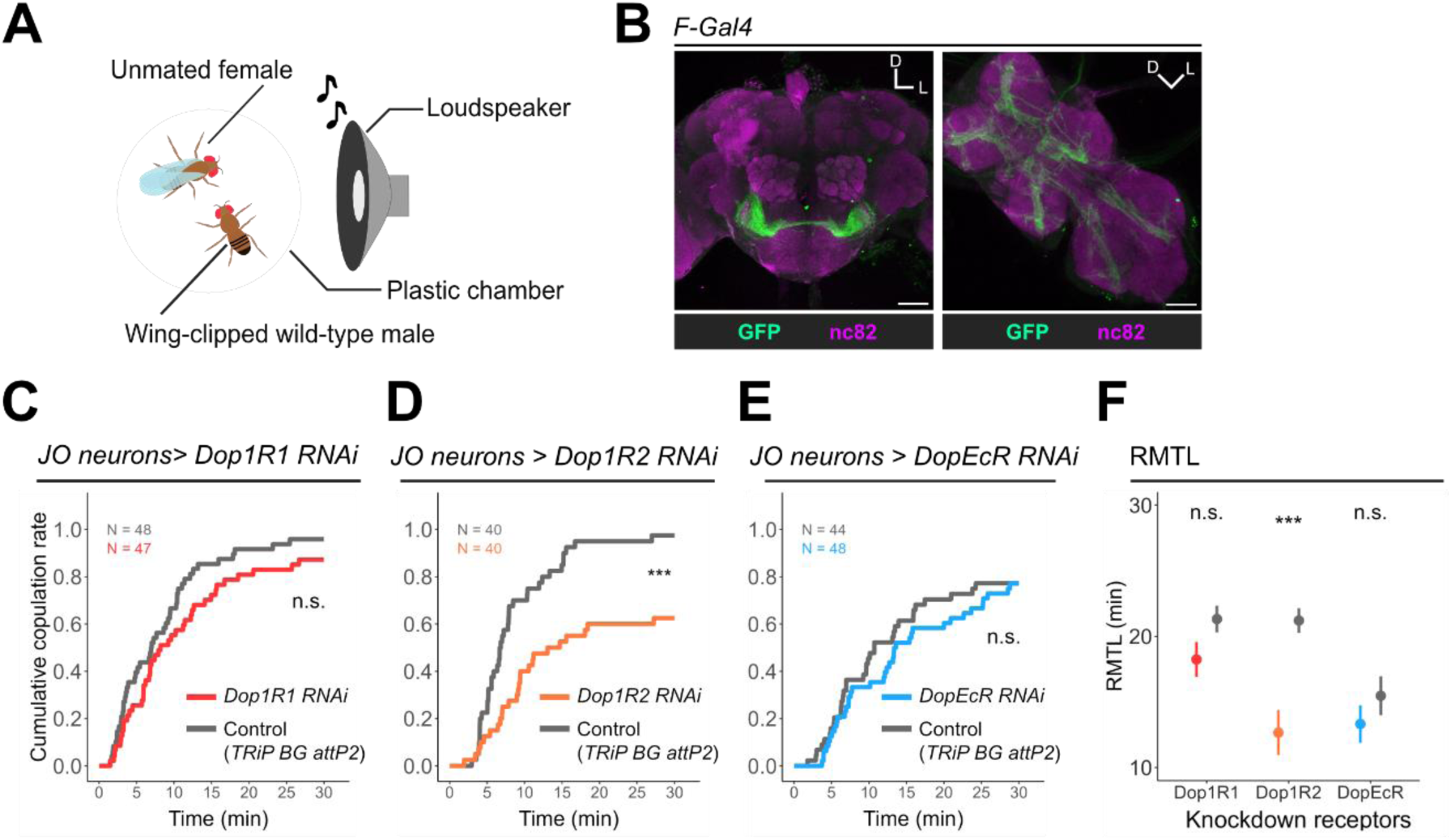
*Dop1R2* knockdown in JO neurons decreases the song response behavior of unmated females. (A) Female copulation assay. An unmated female was paired with a wing-clipped wild-type male in a chamber. An artificial pulse song was played back through a loudspeaker. The experiment was also conducted under a no-sound condition (See Figures S15, S16). (B) *F-Gal4* driver strain was used to express mCD8::GFP markers (green). nc82 antibodies were used for counter-labeling (magenta). Scale bar = 50 µm. (C-E) Cumulative copulation rates during song playback. Females with *Dop1R1* (C), *Dop1R2* (D), or *DopEcR* (E) knockdown in JO neurons were tested. (F) Evaluation of female song response behavior with Restricted mean time lost (RMTL). *F-Gal4* driver strain was used to express each RNAi construct. Dot and vertical line represent the estimated value and standard error, respectively.

Among the three receptor types that are abundantly expressed in JO-AB neurons, only knockdown of *Dop1R2* in JO neurons decreased the song response behavior (Figure 5D), while the knockdown of neither *Dop1R1* nor *DopEcR* affected it (Figures 5C, 5E). Comparison of the overall amount of behavioral responses, represented by restricted mean time lost (RMTL),^13,53^ also detected a significant reduction in song response behavior in the *Dop1R2* knockdown group but not in others, further validating the reduced behavioral responses in *Dop1R2* knockdown females (p = 8.54 × 10^−2^, 2.04 × 10^−5^, and 0.312 for *Dop1R1*, *Dop1R2,* and *DopEcR*, respectively; RMST tests corrected with Benjamini and Hochberg method; Figure 5F; Table 10; See Figures S15, S16 for no-sound controls). Quantifying the walking speeds of *Dop1R2* knockdown and control flies revealed similar activity levels (Figure S17; Table 11), ruling out the possibility that the reduced copulation phenotype observed in the *Dop1R2* knockdown group was due to aberrant locomotor activity in female flies. These analyses, therefore, suggest that the modulation via *Dop1R2* in JO neurons, but not *Dop1R1* and *DopEcR*, affects song response behavior in unmated females.

**Table 10:** Statistical summary of female receptivity using RMTL corrected with Benjamini and Hochberg method.

| Figure | Strain | Sound | RMTL (min) | SE | p-value | Adjusted p-value |
| --- | --- | --- | --- | --- | --- | --- |
| 5C,<br>S16A | *F-Gal4>Dop1R1 RNAi* | 35-ms IPI | 18.2 | 1.32 | 0.0650 | 0.0854 |
|  | *F-Gal4>TRiP* background |  | 21.3 | 1.02 |  |  |
| S15A,<br>S16A | *F-Gal4>Dop1R1 RNAi* | No sound | 0.00532 | 0.00526 | 0.0854 | 0.0854 |
|  | *F-Gal4>TRiP* background |  | 1.01 | 0.583 |  |  |
| 5D,<br>S16B | *F-Gal4>Dop1R2 RNAi* | 35-ms IPI | 12.6 | 1.70 | 1.02E-05 | 2.04E-05 |
|  | *F-Gal4>TRiP* background |  | 21.2 | 0.934 |  |  |
| S15B,<br>S16B | *F-Gal4>Dop1R2 RNAi* | No sound | 0 | 0 | 0.311 | 0.311 |
|  | *F-Gal4>TRiP* background |  | 0.423 | 0.418 |  |  |
| 5E,<br>S16C | *F-Gal4>DopEcR RNAi* | 35-ms IPI | 13.3 | 1.43 | 0.296 | 0.312 |
|  | *F-Gal4>w<sup>1118</sup>* |  | 15.5 | 1.48 |  |  |
| S15C,<br>S16C | *F-Gal4>DopEcR RNAi* | No sound | 0 | 0 | 0.312 | 0.312 |
|  | *F-Gal4>w<sup>1118</sup>* |  | 0.180 | 0.178 |  |  |
| 6A,<br>S16D | *JO15>Dop1R2 RNAi* | 35-ms IPI | 15.0 | 1.76 | 3.82E-06 | 6.57E-06 |
|  | *JO15>TRiP* background |  | 24.1 | 0.892 |  |  |
| S15D,<br>S16D | *JO15>Dop1R2 RNAi* | No sound | 0.168 | 0.166 | 0.575 | 0.575 |
|  | *JO15>TRiP*<br>background |  | 0.417 | 0.413 |  |  |
| 6B,<br>S16E | *R74C10>*<br>*Dop1R2 RNAi* | 35-ms IPI | 20.0 | 1.39 | 0.0192 | 0.0384 |
|  | *R74C10>*<br>*TRiP* background |  | 23.9 | 0.890 |  |  |
| S15E,<br>S16E | *R74C10>*<br>*Dop1R2 RNAi* | No sound | 0.242 | 0.239 | 0.212 | 0.212 |
|  | *R74C10>*<br>*TRiP* background |  | 0.909 | 0.478 |  |  |
| 6C,<br>S16F | *R91G04>*<br>*Dop1R2 RNAi* | 35-ms IPI | 16.3 | 1.70 | 0.00817 | 0.0163 |
|  | *R91G04>*<br>*TRiP* background |  | 22.2 | 1.48 |  |  |
| S15F,<br>S16F | *R91G04>*<br>*Dop1R2 RNAi* | No sound | 1.10 | 0.655 | 0.107 | 0.107 |
|  | *R91G04>*<br>*TRiP* background |  | 3.07 | 1.03 |  |  |

**Table 11:** Statistical summary of locomotor activity using Welch’s t-test.

| Figure | Condition | t-value | p-value | Cohen's d |
| --- | --- | --- | --- | --- |
| S17 | *F-Gal4 > Dop1R2 RNAi*,<br>*TRiP* background | 0.246 | 0.813 | 0.155 |

To validate the contribution of *Dop1R2* expression in JO-AB neurons, we next restricted the knockdown population to these neurons using the *JO15* strain (Figure S14B).^16,54^ *Dop1R2* knockdown in JO-AB neurons reduced song response behavior (Figure 6A), manifested by a significant decrease of RMTL (p = 6.57 × 10^−6^; RMST tests corrected with Benjamini and Hochberg method; Figure 6D; Table 10; See also Figures S15, S16 for no-sound controls). To compare the changes in receptivity levels between the knocked-down populations, we calculated ΔRMTL as the reduction in RMTL value from the control group resulting from *Dop1R2* knockdown.^55,56^ ΔRMTL in JO-AB neurons was similar to that in all subgroups of JO-neurons (mean ΔRMTL = 8.55 and 9.16, respectively), suggesting that JO-AB neurons primarily contribute to the effect of *Dop1R2* expression in JO neurons for this behavioral phenotype (Figures 5F, 6D, 6E; Table 12).

**Figure 6:**
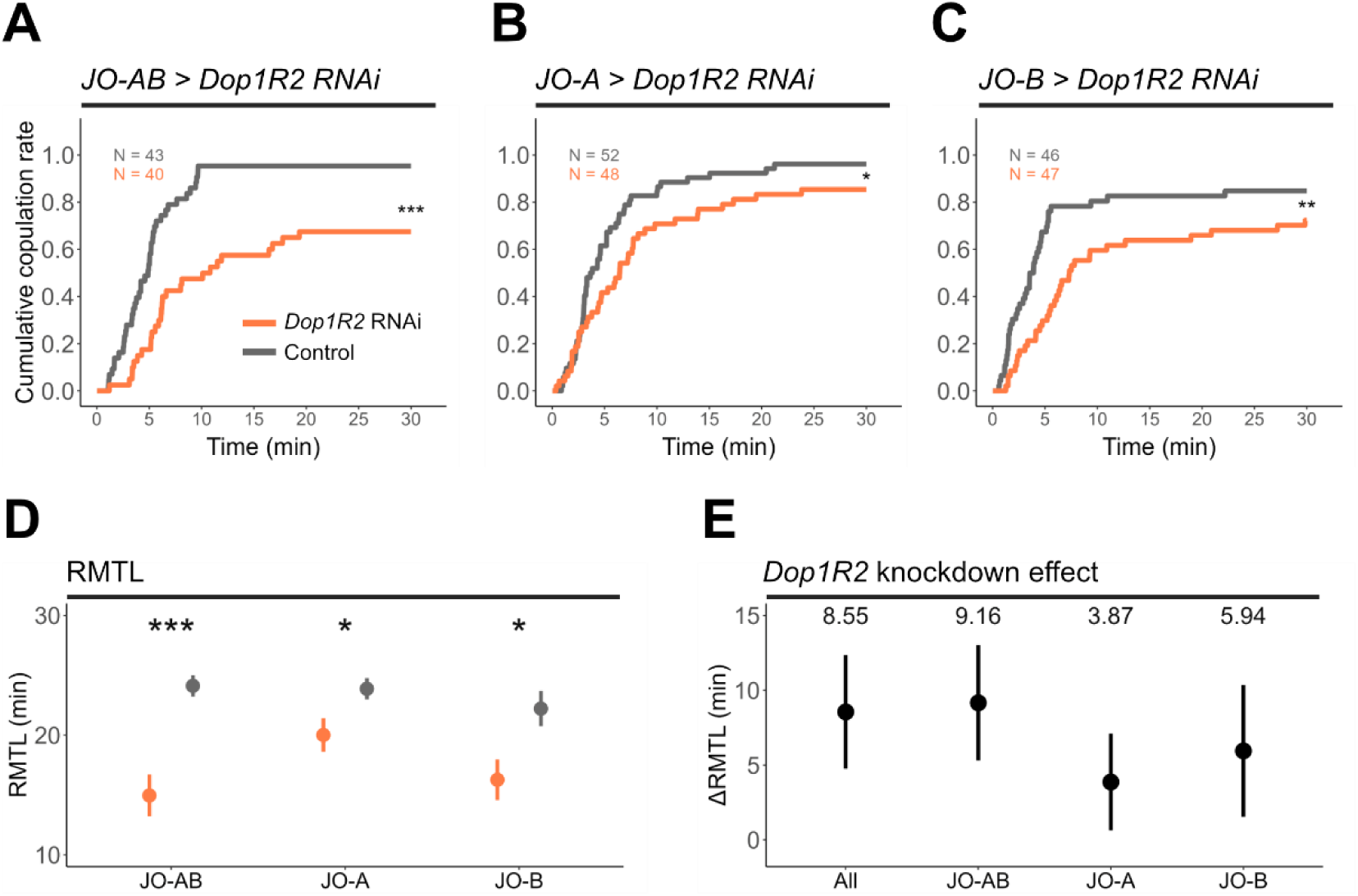
*Dop1R2* knockdown in JO-AB decreases the song response behavior of unmated females. (A-C) Cumulative copulation rates during song playback. Unmated females with *Dop1R2* knockdown were tested. RNAi-mediated knockdown was induced in JO-AB (A), JO-A (B), and JO-B neurons (C). Gal4 driver strains that selectively label JO-AB neurons, JO-A neurons, and JO-B neurons (*JO15*, *R74C10*, and *R91G04* strains, respectively) were used to express the *Dop1R2* RNAi construct. (D) Comparison of female song response behavior using RMTL. Dot and vertical line represent the estimated value and standard error, respectively. (E) *Dop1R2* knockdown effect. ΔRMTL represents the difference in RMTL between experimental and control groups in each condition. ΔRMTLs of *Dop1R2* knockdown in all subgroups of JO neurons, JO-AB neurons, JO-A neurons, and JO-B neurons are shown. Dot and vertical line represent the estimated value and 95% Confidence Interval, respectively.

**Table 12:** Δ RMTL values.

| Figure | Strain | Sound | $\Delta$ RMTL (min) |
| --- | --- | --- | --- |
| 6E | *F-Gal4>Dop1R2 RNAi*, *TRiP* background | IPI35 | 8.55 |
| 6E | *JO15>Dop1R2 RNAi*, *TRiP* background | IPI35 | 9.16 |
| 6E | *R74C10>Dop1R2 RNAi*, *TRiP* background | IPI35 | 3.87 |
| 6E | *R91G04>Dop1R2 RNAi*, *TRiP* background | IPI35 | 5.94 |

Finally, we estimated the contribution of Dop1R2 in each JO subgroup using two GAL4 drivers (Figures S14C, S14D),^19,21^ which selectively label JO-A and JO-B neurons respectively. *Dop1R2* knockdown in either JO-A or JO-B significantly decreased song response behavior (p = 3.84 × 10^−2^ and 1.63 × 10^−2^ for JO-A and JO-B, respectively; RMST tests corrected with Benjamini and Hochberg method; Figures 6B, 6C; Table 10; See Figures S15, S16 for no-sound controls). Notably, knockdown in both subgroups of JO neurons induced a greater reduction in behavior (Figure 6A) than knockdown in single subgroups, as depicted in the ΔRMTL (ΔRMTL = 3.87 min for JO-A and 5.94 min for JO-B; Figures 6D, 6E; Table 12). The summation of ΔRMTLs from these two single-subgroup knockdowns seems close to the calculated *Dop1R2* knockdown effect in JO-AB neurons (Figure 6E, Table 12), suggesting that *Dop1R2* expressed in JO-A and JO-B neurons collaborate to enhance the song response behavior of unmated female flies.

## III. Discussion

Here, we demonstrate a mating state-dependent modulation of auditory processing via the dopaminergic system in the female fruit-fly brain. Three types of dopamine receptors are abundantly expressed in JO-AB neurons, some of which presumably mediate direct synaptic transmission from dopaminergic neurons to JO-AB neuron axons. Among these receptor types, Dop1R2 expression significantly contributed to sound-evoked neuronal responses, by enhancing sound responses of JO-A neurons in unmated females. The enhanced JO-A responses resulting from Dop1R2 expression were not observed in mated females, suggesting a mating-state-dependent dopaminergic effect on neuronal responses. Behavioral analyses further suggested that Dop1R2 expression in JO-A and JO-B neurons additively enhances the song response behavior of unmated female flies, highlighting the contribution of Dop1R2 in both JO neuron subgroups to maximize song response behaviors. Interestingly, JO-A responses in newly eclosed females were not affected by *Dop1R2* knockdown, highlighting mating drive as a key factor for the state-dependent Dop1R2 effect. These findings suggest that female mating drive affects the properties of auditory sensory neurons via dopamine signals, thereby modulating female behavioral responses to the courtship sounds emitted by males for accepting copulation (Figure S13B).

### Dopamine signals to JO-AB neurons

Dopamine is involved in state-dependent sensory processing in various animals. In mice, the decrease of dopamine synthesis in olfactory mucosa during starvation contributes to olfactory enhancement.^24^ In flies, dopamine regulates the sensitivity of gustatory sensory neurons in a hunger-state-dependent manner.^25^ This study suggests that dopamine modulates auditory sensory neurons depending on females’ mating drive in *Drosophila*, highlighting the role of dopamine in reflecting animals’ states to their sensory neurons. Although many dopaminergic neuron clusters have been identified in the *Drosophila* brain (e.g., PAM, PAL, T1, Sb, PPL1, PPL2ab, PPL2c, PPM1, PPM2, and PPM3 clusters),^57^ it is unclear whether any of these clusters affect auditory processing. Several dopaminergic neurons innervating the AMMC, identified in *Drosophila* neuron databases (Tables 4, 5), serve as promising candidates for providing dopaminergic modulations to JO-AB neurons.

Most JO neurons express dopamine receptors, but a limited number receive direct synaptic connections from dopaminergic neurons (Figures 1, 2). In mammals, volume transmission is suggested to be the primary mode in dopamine circuits.^58^ In *Drosophila*, some studies suggest that dopamine diffuses outside of the postsynaptic region, implying that volume transmission of dopamine takes place in addition to direct synaptic transmission.^59,60^ These two modes of dopaminergic transmission potentially coordinate to reflect the internal states of JO neurons. Alternatively, it is also possible that dominant effects occur via volume transmission. Further validation is necessary to understand the synaptic mechanism that mediates this modulation.

### Dop1R2 function in JO-AB neurons

We found that the sound-response of JO-A neurons in unmated females is enhanced by Dop1R2, which is classified as a metabotropic D1-like receptor in fruit flies.^61^ In the vertebrate hearing system, the function of D1 receptors has been analyzed pharmacologically, revealing its effect in enhancing the neural response in the cochlea: Application of a D1 receptor agonist SKF38393 increased the cochlear compound action potential in guinea pigs,^62^ and microphonic potentials and intracellular calcium levels in zebrafish hair cells.^63^ Our findings align with these reports, together highlighting the excitatory role of D1 receptors at the first stage of auditory processing. Notably, Dop1R2 expression levels in JO-A axons were stable before and after mating (Figure S11). Given that L-DOPA feeding did not increase calcium responses in mated females (Figure S12), changes in the downstream pathway of Dop1R2 signaling might mediate the specific dopamine effect in unmated female flies (Figure S13A).

Previous studies have shown that *Drosophila* Dop1R2 activates intracellular cyclic AMP (cAMP) and calcium signaling pathways upon dopamine binding.^31,32,64^ These pathways may increase the response amplitude of JO-A neurons by modulating various cellular processes.^65–67^ Our model-based estimation of the calcium dynamics further suggests that Dop1R2 expression accelerates the intracellular calcium increase (𝑟) and delays its decrease (𝑑_2_) in JO-A neurons (Figure 3), together possibly resulting in a net increase in neurotransmitter releases during the stimulus and facilitation of calcium responses to a subsequent stimulus. Identifying the cellular processes responsible for these kinetic changes would enhance our mechanistic understanding of Dop1R2-mediated modulation in JO-A neurons.

Although *Dop1R2* knockdown in either JO-A or JO-B neurons reduced the song response behavior of females (Figure 6), it did not decrease the sound-evoked calcium responses in JO-B neurons (Figure 3). A possible explanation is that dopamine signals through Dop1R2 affect the molecular machinery involved in neurotransmission in JO-B neurons without affecting the stimulus-induced calcium responses. Monitoring neurotransmitter release directly from JO-B neurons using genetically encoded neurotransmitter sensors, e.g., GRAB sensor for acetylcholine,^68^ would clarify this possibility, further highlighting the distinct modulatory mechanisms regulating JO-A and JO-B auditory functions.

### The role of JO-A neurons during song response behavior

*Dop1R2* knockdown in JO-A neurons reduced female song response behaviors (Figure 6), suggesting that JO-A neurons play a crucial role in this behavior. In male *Drosophila*, TNT-based suppression of JO-A neurons reduced the song-response behavior,^19^ together suggesting their involvement in processing song information in both males and females. However, JO-A neurons are not strongly connected with AMMC-B1 neurons, the main secondary auditory neurons that process song information in the fly brain.^21^ Thus, the neural circuit mechanisms underlying their function in song response behavior remain poorly understood, unlike JO-B neurons which have strong synaptic connections with AMMC-B1 neurons.^21^ Notably, AMMC-A2 neurons, one of the downstream neurons of JO-A neurons,^69,70^ have been identified as pulse-song-preferring neurons.^7^ AMMC-A2 neurons further connect to pC2l and vpoIN neurons in two synaptic steps (i.e., hops),^7^ which are also tuned to the pulse song and involved in regulating mating acceptance/rejection in females.^10,71^ These findings support the idea that AMMC-A2 neurons serve as another type of song-relaying secondary auditory neurons, alongside AMMC-B1 neurons, in song response behavior. Exploring the interaction between the downstream neural circuits of JO-A neurons, possibly including AMMC-A2 neurons, and the song-relay circuit leading to the mating decision of females will elucidate how the two major types of auditory sensory neurons, JO-A and JO-B, coordinate to affect song response behaviors in flies. The different *Dop1R2* knockdown phenotypes in neuronal responses observed between JO-A and JO-B, along with their additive role for Dop1R2-mediated song response behavior of females, are likely related to their distinct roles in song information processing; further study is needed to understand this relationship.

### Possible modulations across multiple circuit levels

In the *Drosophila* olfactory pathway, the mating status of females modulates the neuronal responses to polyamines in both sensory and higher-order neurons.^72^ While the modulation in olfactory sensory neurons occurs only within 24 h after mating, the effect on higher-order neurons lasts for 3 to 5 days. This suggests that different mechanisms underlie these modulations in the olfactory pathway, orchestrating the state-dependent behaviors.^72–74^ It is thus intriguing to test whether responses of higher-order auditory neurons in the song-processing pathway, such as AMMC-B1 neurons and pC2 neurons,^8,9^ are also modulated according to the mating status of female flies, thereby adjusting the properties of the entire song processing pathway.

In the green treefrog, changes in midbrain responses to mating calls after mating specifically occur in the low-frequency range, which covers one of the two key frequency ranges in mating calls that attract females.^1,2^ In contrast, Dop1R2-mediated modulation of JO-A neuronal responses in flies occurs for all sound stimuli tested in this study, suggesting a non-specific dopaminergic effect on JO-A neurons. Previous studies in flies have reported that many higher-order auditory neurons in the song-processing pathway are tuned to the sound stimuli carrying species-specific temporal features (e.g., K. Wang et al.^10^; Yamada et al.^18^), whereas JO-A neurons respond to a broad range of sounds.^19^ Testing whether modulation occurs in higher-order auditory neurons, and if so, whether it happens in a sound-specific manner, may help us better understand the state-dependent modulation of females’ response to courtship songs. Moreover, in addition to the dopaminergic receptors, JO neurons express various types of neurotransmitter receptors (Figure S1A), suggesting that auditory sensory neurons receive multiple neurotransmitters, likely at their axons. How these neurotransmitters modulate JO neuron function is an interesting question for future research.

### Mechanisms underlying mating status-dependent modulations

Neural mechanisms underlying mating status-dependent modulations have been studied in several model animals. In mice, signaling factors in seminal fluid that influence female immune response are transferred from males to females during copulation.^75^ While factors equivalent to the *Drosophila* SP, which significantly affect female behaviors, have not yet been identified,^76^ sex hormones such as estrogen and progesterone, which increase during pregnancy,^77,78^ are suggested to contribute to changes in food intake behaviors in mammals.^79^ In female *Drosophila*, the SP transferred from the male during copulation triggers the post-mating behavioral switch via SP receptors.^48–51^ Neurons modulated by SP alter the gustatory response behaviors of female flies to salt and yeast, suggesting that SP modulates gustatory processing neurons based on mating status.^80^ Additionally, the post-mating increase in polyamine response behaviors is accompanied by an enhanced neuronal response to polyamines, induced by the upregulation of SP receptor expression in chemosensory neurons.^72^ The involvement of a juvenile hormone (JH) in mating status-dependent modulations in olfactory sensory neurons has also been reported,^81^ suggesting that multiple cellular mechanisms coordinate to regulate their sensory behaviors.

This study suggests that SP has a minor effect on the mating status-dependent modulation of auditory sensory neurons in female flies, where the *Dop1R2* knockdown effect on JO-A neuronal responses was absent in both mated and pseudo-unmated females. Other factors such as sperm, seminal fluid, courtship signals from males, and/or mechanical stimulation during mating potentially link the mating status to the suppression of the Dop1R2 signaling pathway in mated females. Interestingly, our analysis of the single-nucleus RNA-seq dataset suggests the expression of JH receptors in putative JO-AB neurons (Figure S1B), implying that the JH pathway functions in these neurons. Understanding whether the JH systems influence Dop1R2 signaling in JO-AB neurons requires further investigation.

Although mating-status-dependent auditory modulations have been observed in several animal species, the underlying neural mechanisms have yet to be well studied compared to the post-mating modulation of olfactory and gustatory processing. A possible reason may be the paucity of identified phenomena in model animals where molecular genetic tools to monitor and manipulate neuronal functions are well-developed. This study finds that auditory processing is modulated based on the mating state of *Drosophila*, opening avenues to explore the underlying molecular mechanisms.

### Modulations across different sensory modalities

Mating status-dependent modulation of sensory processing in females has been observed in various sensory modalities (Drummond-Barbosa & Spradling^82^; Hussain et al.^72^; Walker et al.^80^ for flies; Choo et al.^83^ for mice; Duffy et al.^84^; Kölble et al.^85^ for humans in olfaction and gustation). Although numerous studies have focused on chemosensory systems, such as olfaction and gustation, a few studies have reported modulations in the visual and auditory systems. One such example is found in cichlids (*Astatotilapia burtoni*), where reproductive females exhibit enhanced visual sensitivity and stronger behavioral responses to male courtship displays compared to non-reproductive females.^86^ Another example is crickets (*Teleogryllus oceanicus*), where the behavioral latency of females to the male’s calling song becomes longer after mating.^87^ These modulations likely enhance fitness by improving the females’ ability to detect males during periods when they can produce offspring. Given that animal behaviors heavily depend on multimodal sensory integration, coordinated modulation across different sensory modalities is likely essential to support behavioral changes associated with mating states. The dopaminergic system likely plays a pivotal role in this cross-modal regulation, facilitating adaptive behaviors.

### Limitations of the study

Our research revealed that Dop1R2 is involved in mating-status dependent modulation in auditory sensory neurons in *Drosophila*. However, this study has the following limitations: (1) The ratio of Dop1R1-positive cells differs between the RNA-seq data and the expression patterns of T2A-Gal4 knock-in driver lines. This discrepancy may stem from differences between transcriptional (RNA-seq) and translational (marker expression) levels. It may also be influenced by experimental conditions (e.g., sex, rearing environment), as the RNA-seq data was obtained from a mixed population of 5-day-old males and females,^23^ while the marker-expression patterns in this study were observed exclusively in 5-10 days old females. The expression level of Dop1R1 may be dynamically regulated by these factors, but further investigation is needed to clarify this. (2) It is reported that *trans*-Tango system has caveats, leading to false positives and negatives.^39,88^ Although we followed the protocol to suppress the false negatives by raising flies at 18°C, we cannot exclude the possible detection of false signals. (3) DopEcR is highly expressed in JO neurons; however, RNAi-mediated knockdown did not result in any detectable phenotypic changes. This suggests the possibility of synergistic or antagonistic interactions between DopEcR and Dop1R2. To explore these potential interactions, further studies using double-knockout experiments are warranted. (4) Theoretically, GFP_1-10_ is expressed excessively due to the amplification effect of the *Gal4/UAS* system. The limiting factor for reconstituted GFP expression is the level of the GFP_11_ fragment, which reflects the endogenous expression of Dop1R2. Therefore, Dop1R2 expression can be quantified by measuring GFP fluorescence. However, to our knowledge, there is no prior publication validating the sensitivity of the split-GFP method in detecting expression level differences.

## STAR Methods

### Key resources table

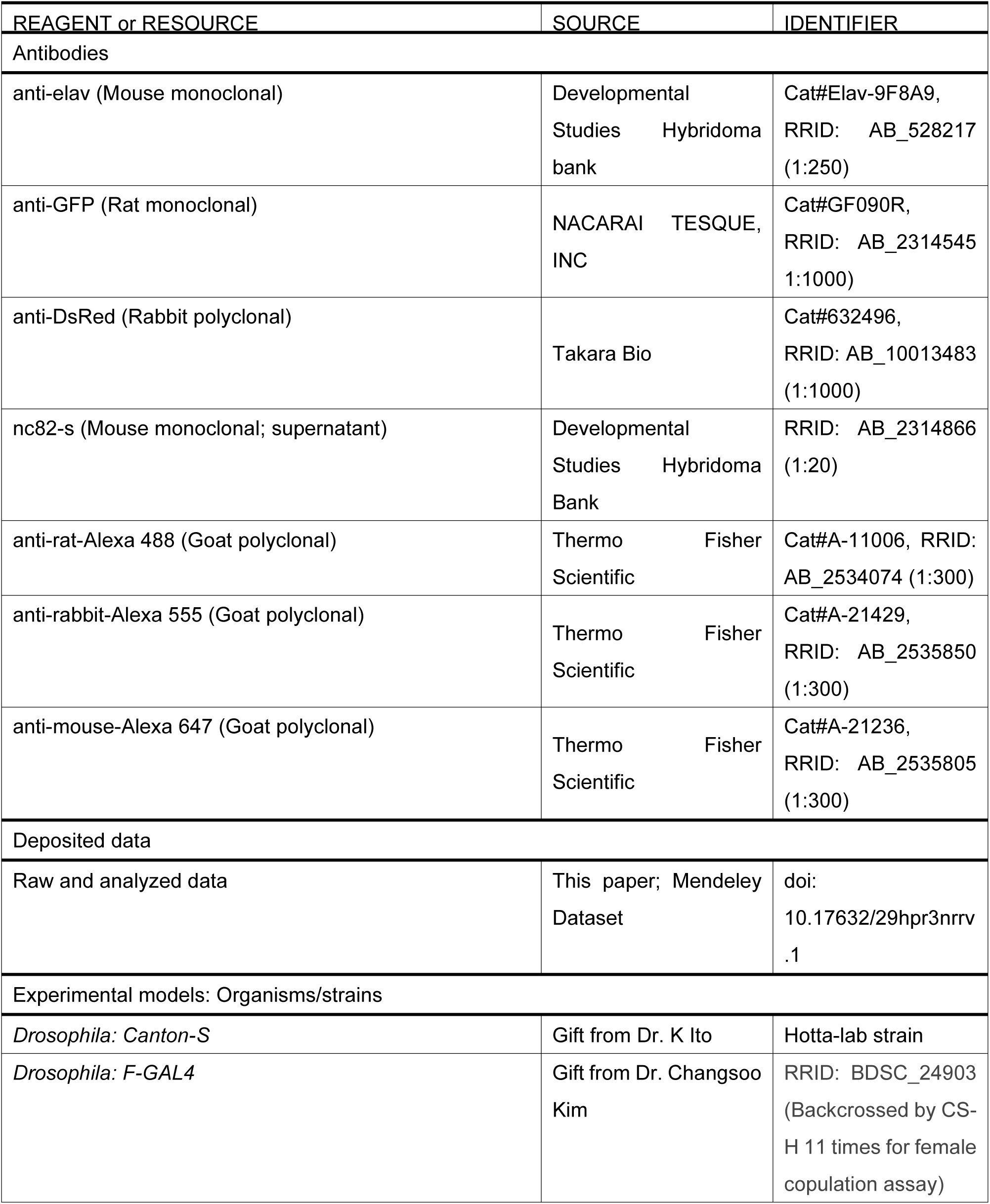
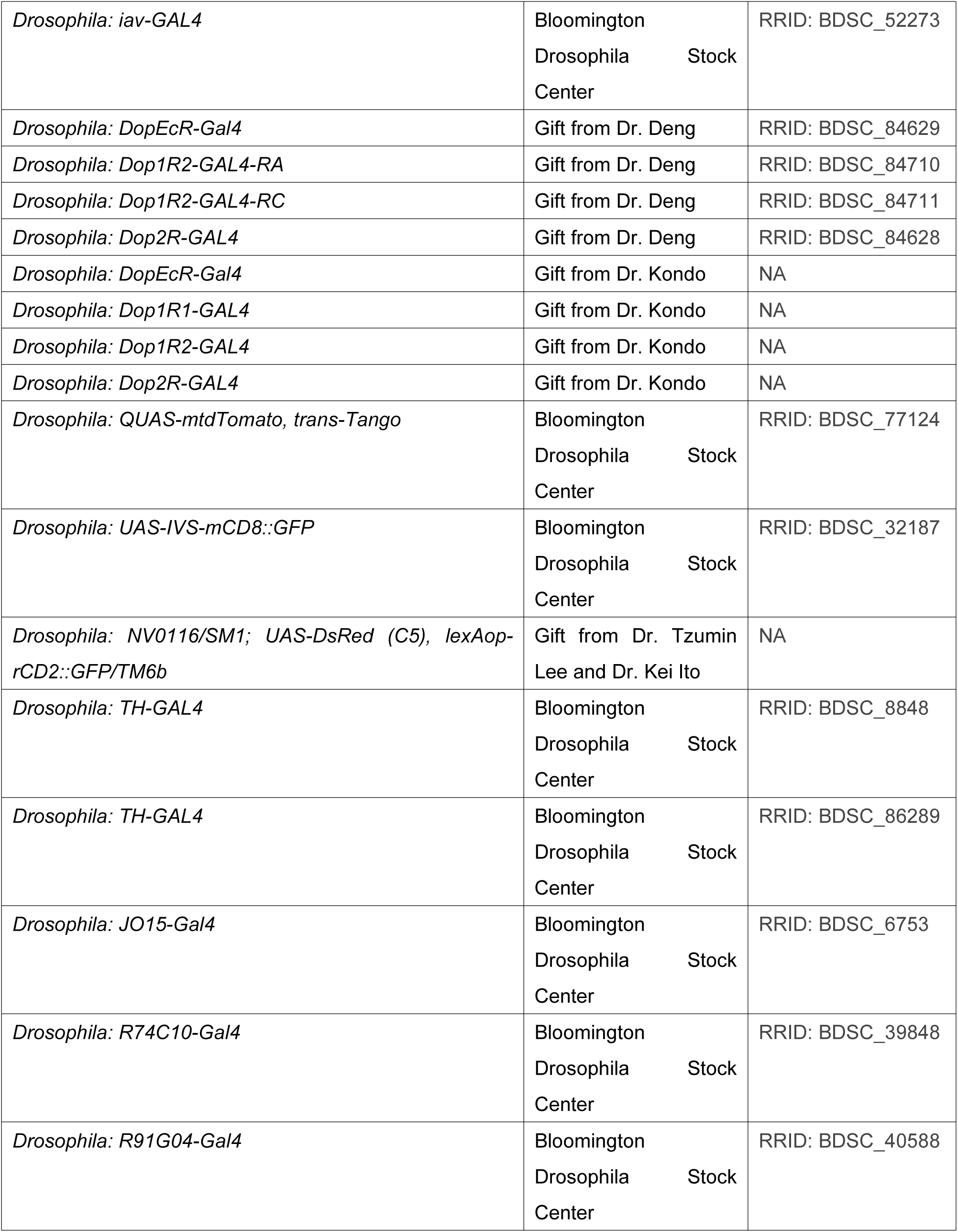
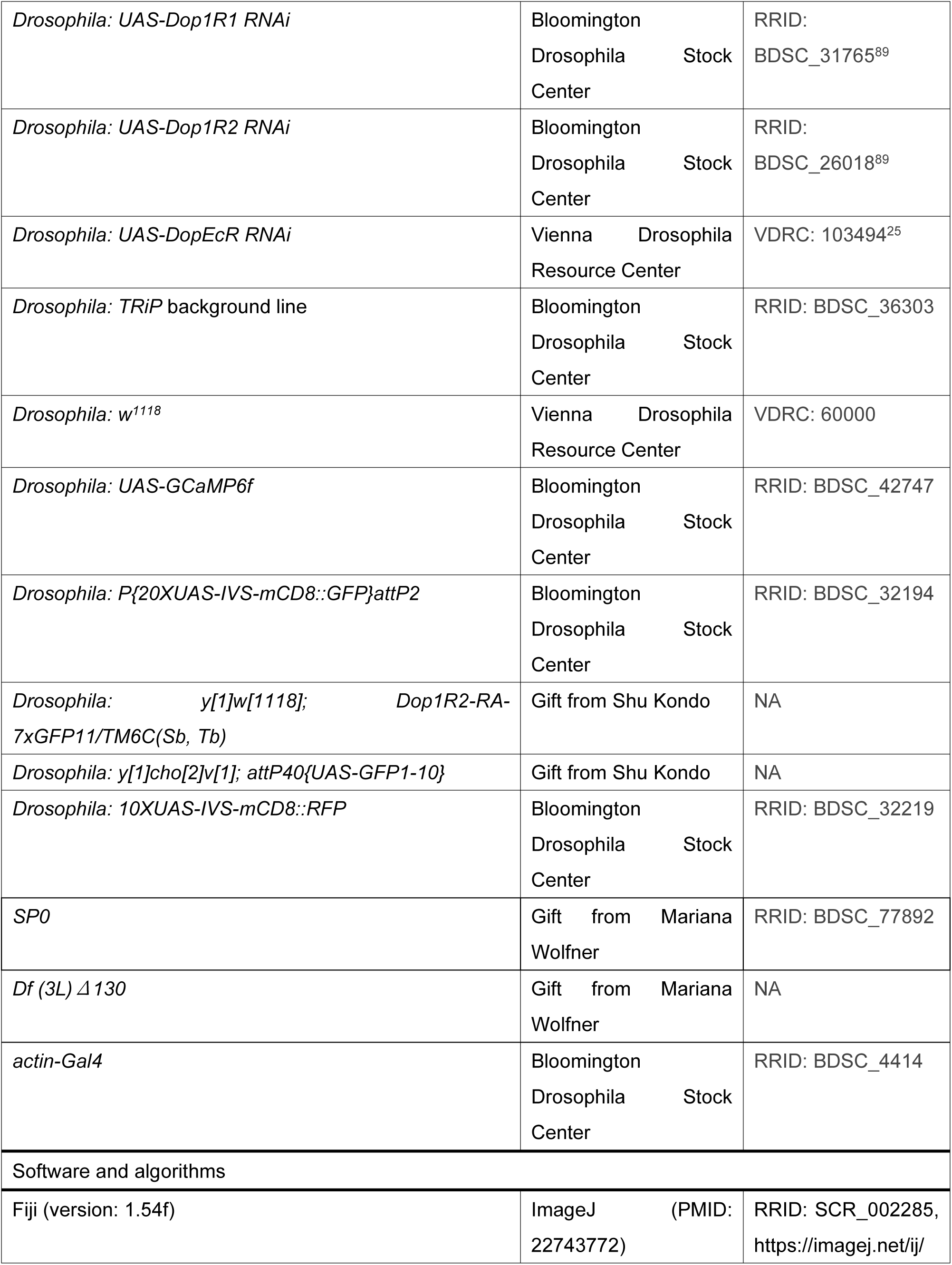
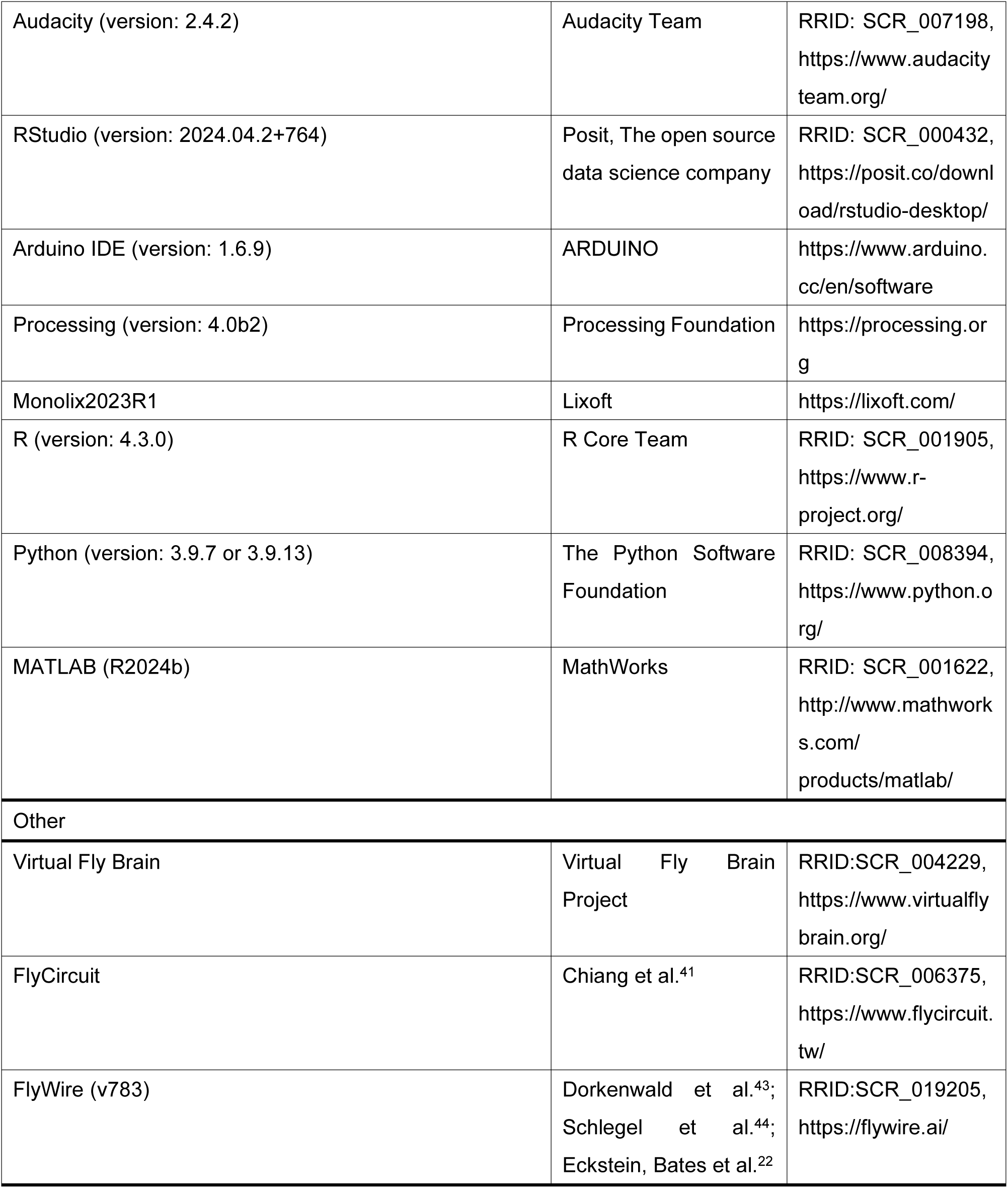

### Experimental Animals

Fruit flies were raised on yeast-based food at 25°C and 40–60% relative humidity under a 12 h light/12 h dark (12 h L/D) cycle unless otherwise noted. Transgenic fly strains were used for the *Gal4/UAS*^90^ and *LexA/lexAop*^91,92^ systems. *Canton-S* was used as a wild-type *D. melanogaster* strain. For the transsynaptic analysis (Figure 2), we used two types of Gal4 driver strains: (1) promoter-Gal4 line, in which a promoter-Gal4 construct was inserted into the genome using transposase (Figure S4A),^40^ and (2) a knock-in line where Gal4 was fused to the C-terminus of the endogenous tyrosine hydroxylase (TH) protein (Figure S4B).^35^ Detailed information on the *Drosophila* strains is described in the Key Resources Table. Genotypes of flies used for each experiment are listed in Table S1.

To prepare unmated and mated females for calcium imaging experiments and split-GFP analysis, adult females collected within 8 h after eclosion were raised in groups of 3 to 10. Unmated females were kept in vials for 5-7 days before being used for the experiments. Mated and pseudo-mated females were kept in vials for 4-6 days, after which three females each were transferred to a food vial containing 3-10 unmated wild-type or SP null (*SP0*/ *Df(3L)Δ130*) males, where they were kept for 12-24 h. This procedure resulted in fertilization of over 98% of females, as confirmed by quantifying the population of females that produced larvae after mating with wild-type males. To prepare newly eclosed females, adult females collected within 6 h after eclosion were raised in groups of 3 to 10 for 20-30 h.

For the L-DOPA feeding experiment, adult females collected within 8 h after eclosion were raised in groups of 3 to 10 for 3-5 days. Females were starved for 15-18 h by placing in a vial containing a paper towel moistened with water. The females were then paired with wild-type males as described above and placed in a vial containing food with L-DOPA (0.3% (w/v) L-DOPA (3-(3,4-Dihydroxyphenyl)-L-alanine, #D0600, Tokyo Chemical Industry Co., Ltd), 0.073% (w/v) HCl (#37314-15, NACALAI TESQUE, INC.), 9% (w/v) glucose (Glul Final, Sanei Toka Co., Ltd), 5% (w/v) Extract of yeast (#103753, Merck KGaA) and 1% (w/v) agarose (#312-01193, NIPPON GENE CO., LTD.) in water). For controls, food without L-DOPA was used. Flies were kept for 12-24 h and used for calcium imaging.

At the time of calcium imaging, unmated and mated females were 5-7 days old after eclosion, while newly eclosed females were 26-36 h after eclosion.

### Immunohistochemistry

Immunohistochemistry was performed as described previously with minor modifications.^18^ Adult females 5-10 days after eclosion were ice-anesthetized, washed in 99.5 % ethanol (14033-81, KANTO CHEMICAL CO., INC.) for about 30 sec to remove the oil on the body surface, and dissected in phosphate-buffered saline to obtain brains and antennae (PBS; #T9181, Takara). After dissection, these tissues were fixed for 1 h in 4% paraformaldehyde/PBS (163-20145, Wako Pure Chemical Industries, Ltd) on ice or 30 min in glyoxal solution (3% (w/v) formaldehyde (#28908, Thermo Fisher Scientific), 1% (w/v) glyoxal (#077-05291, FUJIFILM Wako Pure Chemical Corporation), 0.1% (w/v) methanol (#34885, Sigma-Aldrich) in PBS at room temperature.^93,94^ Samples were then washed twice with PBS containing 0.5% Triton X-100 (PBT) (#X100-500ML, Sigma-Aldrich), and incubated overnight in PBT at 4°C. Subsequently, samples were incubated with primary and secondary antibodies for three days each at 4°C. The antibodies are listed in Key Resources Table. After washing the samples with 0.5% PBT, samples were transparentized in 50% and 80 % Glycerol (075-04751, Wako Pure Chemical Industries, Ltd) sequentially.

### Confocal microscopy and image processing

Serial optical sections of brains and antennae were obtained at 0.50 to 0.84 µm intervals with a resolution of 512 × 512 pixels using a FLUOVIEW FV1200, FV1000D, or FV3000 laser-scanning confocal microscope (Olympus) equipped with a silicone-oil-immersion lens UPLSAPO 30xS (NA = 1.05; Olympus) or UPLSAPO 60xS (NA = 1.30; Olympus). Image size, brightness, and contrast were adjusted using Fiji (version: 2.0.0-rc-69/1.52p).^95^

### Analysis of expression of T2A-Gal4 knock-in driver lines in the antenna

To quantify the ratio of Gal4-positive JO neurons, we randomly extracted three optical slices from each serial-image dataset of JO. In each section, we counted the number of the *Gal4* > tdTomato signals overlapping with Elav signals to examine the percentage of JO neurons expressing Gal4 signals using the “Cell Counter” plugin in Fiji (version: 2.0.0-rc-69/1.52p).

### Gene expression analyses using single-nucleus RNA-seq data

Fly Cell Atlas (www.flycellatlas.org), a platform for single-nucleus RNA sequence dataset of fruit flies,^23^ was used for gene expression analyses. To estimate the proportion of JO neurons that express neurotransmitter receptors, the antennal dataset (antenna.h5ad; collected from male and female antennae)^23^ was loaded using SeuratDisk (ver. 0.0.0.9020) for R (ver. 4.3.0). The cell cluster previously annotated as “Johnston Organ Neuron”^23^ was extracted using the subset function of the Seurat package (ver. 4.3.0.1).

The proportion of JO neurons that express each dopamine receptor was calculated using the WhichCells function of Seurat package (ver. 4.3.0.1). To estimate the expression level of SP and JH receptors in JO-AB neurons, we extracted Dop1R2-expressing cells from annotated “Johnston Organ Neuron”^23^ as Dop1R2-T2A-Gal4 signals are exclusively expressed in JO-AB neurons (Figure 1). Following a previous study,^23^ the normalized expression level was calculated as follows:

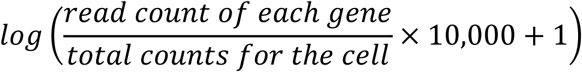

#### *trans*-Tango analysis

For *trans*-Tango analysis, we raised flies for 13-15 days after eclosion at 18°C in accordance with a previous study reporting that rearing at 18°C typically yields the best S/N ratio for *trans*-Tango signals.^39^ The number of *TH>trans-Tango* positive JO neurons was then quantified by counting tdTomato signals that represent *TH-Gal4* > *trans*-Tango signals in JO neuron somata across all confocal optical sections through the second antennal segment using the “Cell Counter” plugin in Fiji (version: 2.0.0-rc-69/1.52p). Signals in the external sensory neurons, located underneath the antennal external sensory bristles, were excluded from the count. To validate that the *trans*-Tango signals in the AMMC were above the autofluorescence level, we acquired confocal images with the same scan settings for the laser power and the detector sensitivity across samples and genotypes (Figure S4G).

### Detecting dopaminergic neurons innervating the AMMC

To extract dopaminergic neurons innervating the AMMC, we used both EM and LM database, Flywire, FlyCircuit, and Virtual Fly Brain.^22,41–44,96^ In the EM database, we searched for dopaminergic neurons using codex, the explorer of the Flywire brain dataset (v783), with the following conditions: input_neuropils == AMMC_L or AMMC_R, nt_type == DA. In the LM database, we searched for dopaminergic neurons using Text-based Search in Flycircuit with the following conditions: Driver == TH-Gal4, Innervation sites == AMMC or ammc. From the hit neurons, we extracted dopaminergic neurons innervating AMMC zone A or zone B. We also observed the morphology of the neurons in Virtual Fly Brain to validate this extraction.

### RT-qPCR

Adult females 5-7 days after eclosion were ice-anesthetized, washed in 99.5 % ethanol (#14032-08, KANTO CHEMICAL CO., INC.) for about 30 sec to remove the oil on the body surface, and dissected in RNA later (#R0901 Sigma-Aldrich) with whole head tissue being collected. Dissected samples were homogenized (Motorized tissue grinder, #12-141-361, Fisher Scientific) in RNAiso (#9109, Takara Bio Inc.). Additional RNAiso was added to the lysed samples to make up a final volume of 1 mL. 200 µL chloroform (#07278-00, KANTO CHEMICAL CO., INC.) was added to each sample and mixed well before incubating at room temperature for 2-15 min. Samples were then centrifuged at 12,000 x g for 15 min at 4°C before the addition of 0.5 mL 2-propanol (#32435-00, KANTO CHEMICAL CO., INC.). Samples were stored in -20°C for 1 hour followed by centrifugation at 12,000 x g for 10 min at 4°C, the supernatant was discarded, and RNA pellet was washed with 0.5 ml of 99.5 % ethanol. Samples were centrifuged at 12,000 x g at 4°C for 5 min. RNA pellet was then washed with 0.5 ml of 75% ethanol and then centrifuged again at 12,000 x g at 4°C for 5 min before the RNA pellet was dissolved in Nuclease Free Water (ReverTra AceTMqPCR RT Master Mix with gDNA Remover, FSQ-301, TOYOBO). RNA sample quality was confirmed using a Nanodrop and stored in -80°C until use. RNA was reverse transcribed (ReverTra AceTMqPCR RT Master Mix with gDNA Remover, FSQ-301, TOYOBO) prior to conducting qPCR (THUNDERBIRDTMSYBR®qPCR Mix, QPS-201, TOYOBO). The housekeeping gene ribosomal protein 49 (*RP49*) was used as the internal control. Dopamine receptor primers were designed using the Primer BLAST (Table S3). For each qPCR 96-well plate (Bio-Rad), three technical repeats for each sample and primer were tested, with the median of these technical values being used for analyses. In total, five biological repeats were conducted.

### Sound file preparation

Artificial pulse songs and 100-Hz pure tone were generated using Audacity (The Audacity Team, version 2.0). For artificial pulse songs, we generated two song types, carrying IPIs of 35 ms and 75 ms, respectively. Both songs are comprised of pulses with a carrier frequency of 167 Hz.^18^ Sound amplitude was adjusted using NR23158-000 microphone (Knowles, NR Series) to 11.1-13.1 mm/s (for pulse songs) or 15.5-18.4 mm/s (for 100-Hz pure tone) particle velocity for the Ca^2+^ imaging experiment and 8.08 mm/s particle velocity for the copulation assay at the peak-to-peak amplitude. For the calcium imaging of the L-DOPA fed females, the sound amplitudes for both pulse songs and 100-Hz pure tone were adjusted to 4.6 and 9.2 mm/s.

### Ca^2+^ imaging - data acquisition

Calcium imaging was performed on female flies as described previously with minor modifications.^13,18^ *F-Gal4* and *iav-Gal4*, fly strains that label most JO neurons spanning all subgroups, were combined to express a Ca^2+^ indicator GCaMP6f^97^ and each dopamine receptor*-*knockdown construct simultaneously. A group of 3 to 10 female flies expressing both transgenes were collected in a food vial within 8 h after eclosion and maintained at 25°C and 40–60% relative humidity under a 12 h L/D cycle until the experiment was conducted. For the imaging, unmated or mated flies five to seven days after eclosion were used; they were anesthetized on ice and stabilized ventral side up onto an imaging plate using silicon grease (SH 44M, Toray, Tokyo, Japan). The mouthparts were removed from the fly head using fine forceps to open a window to monitor GCaMP fluorescence from the brain. A drop of a saline solution (108 mM NaCl, 5 mM KCl, 2 mM CaCl_2_, 8.2 mM MgCl_2_, 4 mM NaHCO_3_, 1 mM NaH_2_PO_4_, 5 mM trehalose, 10 mM sucrose, and 5 mM HEPES, pH 7.5, 265 mOsm)^98^ was added to prevent dehydration. A 35 mm dish (35-mm lummox-film bottom dish, SARDTEDTAG & Co) with a hole at the bottom was set on the window opened at the fly mouthparts and filled with 2 mL of saline solution.

GCaMP fluorescence was detected using a fluorescent microscope (Axio Imager.A2, Carl Zeiss, Oberkochen, Germany) equipped with a water-immersion 20× objective lens (W N-Achroplan, numerical aperture = 0.5; Carl Zeiss), a spinning disk confocal head CSU-W1 (Yokogawa, Tokyo, Japan), a dichroic mirror (405/488/561/640 Di01-T405/488/568/647-13x15x0.5, Semrock, Rochester, NY), and a bandpass filter (FF01-528/38-25, Semrock). An OBIS 488 LS laser (Coherent Technologies, California, US) was used to excite GCaMP6f at 488 nm. The fluorescent images were captured at 10 fps with an exposure time of 100 ms for 10 s using an EM-CCD camera (iXon Ultra 897; Oxford Instruments, Tokyo, Japan) at a resolution of 512 × 512 pixels in water-cooled mode. The imaging setup was operated using Micro-Manager 2.0 (version 2.0).

For sound stimulation, a loudspeaker (Fostex, FF225WK; Foster Electric) equipped with an amplifier (TB10A; Fosi Audio) was located 11 cm from the antenna of the fly. The sound files were played through Arduino Uno R3, using the Adafruit Wave Shield for Arduino Kit v1.1. The onset of fluorescent image recording and sound playback were synchronized using Processing (version: 4.0b2), Arduino IDE (version: 1.6.9), and Arduino UNO R3. Sound playback began 2 seconds after the onset of calcium imaging.

Each individual was randomly presented with the following combinations of sounds four times: 100 Hz, 35 ms IPI and 75 ms IPI, each for 2 seconds. 100-Hz pure tone served as a control sound as it evokes strong responses in both JO-A and JO-B neurons.^45^

### Ca^2+^ imaging - data analysis

Imaging data were analyzed using Fiji (version: 2.0.0-rc-69/1.52p) and R (version 4.2.1) as described previously with minor modifications.^13,18^ Brain movements were corrected using a custom Python script (version: 3.9.7)^99^ or the NormCorre algorithm^100^ with MATLAB, which was manually validated by reviewing all the corrected images. To measure fluorescence intensity, regions of interest (ROIs) were defined using Fiji (version: 2.0.0-rc-69/1.52p). ROIs were placed to cover the axonal region of either JO-A or JO-B neurons.

To quantify the calcium responses, the relative fluorescence change (Δ𝐹/𝐹) of GCaMP6f was normalized as follows: Δ𝐹/𝐹 = (𝐹_𝑛_ − 𝐹_𝐵𝑎𝑠𝑒_)/𝐹_𝐵𝑎𝑠𝑒_, where the mean fluorescence intensity within an ROI at each time point n (𝐹_𝑛_) was divided by the mean fluorescence intensity of the Ca^2+^ response during the 1.9 s preceding the stimulus onset (𝐹_𝐵𝑎𝑠𝑒_).^18^ A three-frame moving average of the Δ𝐹/𝐹 values was used for the time trace graphs and analysis. To correct for fluorescence bleaching, fluorescence from frames 1 to 19 and frames 61 to 100 was used to fit a linear model using the stats package (version 4.3.2) in R.^13^ Fluorescence intensity was corrected by subtracting its fitting function.

### Modeling calcium response dynamics in sound stimulus

We developed the following mathematical model to describe the dynamics of calcium response before (𝑡 < 2), during (2 ≤ 𝑡 ≤ 4), and after (4 ≤ 𝑡) sound stimulation, starting from the onset of calcium imaging (𝑡 = 0):

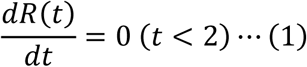

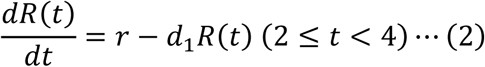

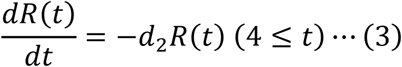

where the sound stimulation started at 𝑡 = 2 . The variable 𝑅(𝑡) is the normalized GCaMP6f fluorescence value (Δ𝐹/𝐹) at time 𝑡 in each fly. The parameters 𝑟, 𝑑_1_, and 𝑑_2_ are the increase rate of calcium response during the stimulus and the decrease rates of calcium response during and after the stimulus, respectively. To estimate these parameters in each fly, we used a nonlinear mixed-effect modeling approach, which includes fixed effect (constant across flies) and random effect (different in flies) for all parameters to consider variability of the responses among flies. Each individual parameter of fly 𝑖, 𝜃_𝑖_(= 𝜃 × 𝑒^𝜋𝑖^), is decomposed as fixed effect (𝜃) and random effect (𝑒^𝜋𝑖^, where 𝜋_𝑖_ is assumed to be drawn from a normal distribution, 𝒩(0, Ω). Parameters of fixed effect and the standard deviation of random effects, 𝜃 and Ω respectively, were estimated with the stochastic approximation Expectation/Maximization (SAEM) algorithm. Then, parameters of individual flies considering random effects were estimated as the mode of their conditional distributions using the empirical Bayes method. Monolix 2023R1 [Monolix 2023R1, Lixoft SAS, a Simulations Plus company] was used as a tool for maximum likelihood estimation applying a nonlinear mixed-effect model to fit the calcium response data.

### Quantification of Split-GFP images – image acquisition

Groups of 3 to 10 female flies were collected in each food vial within 8 h after eclosion and maintained at 25°C and 40–60% relative humidity under a 12 h L/D cycle for seven days. Brains were mounted on a glass slide immediately after dissection, and incubated in SeeDB2G^101^ for at least 30 min to increase transparency. Serial optical sections of brains were then obtained using a FLUOVIEW FV1200 laser-scanning confocal microscope (Olympus) equipped with a silicone-oil-immersion lens UPLSAPO 30xS (NA = 1.05; Olympus) at 0.84 um intervals with a resolution of 512 × 512 pixels. The same scan settings for the laser power and the detector sensitivity (HV; high voltage) were used across samples (GFP, Laser power: 2.6%, HV: 560; RFP, Laser power: 0.7%, HV: 560).

### Quantification of Split-GFP images – analysis

For quantification, one side of the AMMC was used for each individual. The confocal optical images were cropped to 300 × 300 pixels around the AMMC using Fiji (version: 2.0.0-rc-69/1.52p) and Python (version: 3.9.13). To extract the axonal region of JO-A neurons, we binarized RFP images using Imbinarize function for MATLAB. The threshold set by Otsu’s method^102^ in a randomly selected sample was applied to all samples. The GFP intensity was divided by the RFP intensity for normalization in each pixel. Normalized GFP intensity values were averaged in each individual and used for statistical analysis.

### Female copulation assay

Female song response behavior was evaluated using a copulation assay.^18^ *F-Gal4*^52^, *JO15-Gal4*^54^, *R74C10-Gal4*^19^, and *R91G04-Gal4*^21^ were used as driver lines to knockdown the expression of each dopamine receptor in all subgroups of JO neurons, JO- AB neurons, JO-A neurons, and JO-B neurons, respectively (Key Resources Table and Table S1). Wild-type males and genetically manipulated females (knockdown and control groups) were used. Both sexes of flies were collected under ice anesthesia within 8 h after eclosion. Males were wing-clipped and kept in groups of 3 to 10 until experiments were conducted. Females were kept in groups of 3 to 10. These flies were maintained at 25°C and 40-60% relative humidity under a 12 h L/D cycle. Males and females five to seven days after eclosion were used for the assay.

The assays were conducted at 25°C and 40-60% relative humidity during zeitgeber time = 0 to 4. A pair of male and female was gently aspirated into a chamber (1.5 cm diameter, 3 mm deep) without anesthesia. Fly behaviors were recorded for 35 min at 15 fps with a web camera (Logiteck Logicool ®, model: HD Webcam C270). During recording, the artificial courtship song carrying a 35-ms IPI were played back from a loudspeaker (Datio Voice AR-10N, Tokyo Cone Paper MFG. Co. Ltd) via digital amplifier (Lepy LP-2020A; Lepy LP-2024A; Lepy LP-V3S), in which a cycle of one second of pulse sound followed by two seconds silence was repeatedly broadcasted for the 35-min observation period.^18^ The assays were also performed without sound playback as the no-sound condition.

After the assay, the copulation onset in each pair was identified manually. Copulation was judged based on the male being on top of the female for at least five min and the female decreasing her spontaneous behavior.^18,103^

### Locomotor activity analysis

To analyze the locomotor activities of *F-Gal4>Dop1R2 RNAi* females and their control flies, data from the top five pairs with the earliest mating initiation times in each group were used. We tracked the positions of the females during the initial phase of the copulation assay. Using the “Manual Tracking” plugin in Fiji (version: 2.0.0-rc-69/1.52p), we traced their positions over 300 frames following the moment when males oriented and began chasing the females. Total walking distances and walking speeds were calculated, and a ten-frame moving average of the walking speed was applied for plotting.

### Statistical analysis

For all qPCR experiments, median *RP49* values were used to calculate DeltaCt (ΔCt) values for each gene. DeltaDeltaCt (ΔΔCt) values were then calculated within a repeat using control (*actin-Gal4 > TRiP BG attP2* for *Dop1R1* and *Dop1R2*; *actin-Gal4 > w*^1118^ for *DopEcR*) samples as references; all control ΔΔCt values were thus equal to 0. -ΔΔCt to the power of 2 was used for the values in Figure S6. For the copulation assay, RMTL^13^ with the Benjamini and Hochberg method were used to compare the song response behavior between control and experimental groups in each sound condition. For the calcium imaging, Welch’s t-test with the Benjamini and Hochberg method was performed to compare peak calcium responses, increase rates, and decrease rates. For the split-GFP analysis, Welch’s t-test was performed to compare normalized GFP pixel values. stats package (version 4.3.2) and survRM2 package (version 2.2.1) for R (version 4.3.0) was used in these analyses. Cohens’d was calculated using the effsize package (version 0.8.1).

## Acknowledgments

We thank Dr. Yoichi Oda, Dr. Matthew P. Su, Dr. Hiroshi Ishimoto, Dr. Ryo Hoshino, Tai-Ting Lee, and Dr Toshiharu Ichinose for discussions; Dr. Takuro S Ohashi, Daisuke Takaichi and Dr. Ryosuke F. Takeuchi for Python and MATLAB scripts; Dr. Shu Kondo, Dr. Hiromu Tanimoto, Dr. Shun Hiramatsu, Dr. Kuniaki Saito, Dr. Ryusuke Niwa, Drosophila Stock Center, and Vienna Drosophila Resource Center for providing fly strains; Developmental Studies Hybridoma Bank for antibodies.

This study was supported by MEXT KAKENHI Grants-in-Aid for Scientific Research (B) (Grant JP20H03355 to AK), Grant-in-Aid for Transformative Research Areas (A) “iPlasticity” (Grant JP23H04228 to AK), Grant-in-Aid for Transformative Research Areas (A) Hierarchical Bio-Navigation (Grant JP22H05650 to RT) and Materia-Mind (Grant JP24H02200 to AK), Grant-in-Aid for JSPS Fellows (Grant JP24KJ1282 to HY), and JST FOREST (Grant JPMJFR2147 to AK), Japan.

## Data, Materials, and Software Availability

Original data and scripts have been deposited at Mendeley Data: (https://data.mendeley.com/preview/29hpr3nrrv?a=3fb6337d-0680-4893-84fc-9d1ffe549397).

## Author contributions

Conceptualization, H.Y, M.H., and A.K.; Methodology, H.Y., M.H., S.Y., S. Iwanami, S. Iwami, R.T., Y.I., and A.K.; Investigation, H.Y., M.H., and A.K.; Resources, A.K.; Writing,

H.Y. and A.K.; Funding Acquisition, H.Y., R.T., and A.K.; Supervision, R.T., Y.I., and A.K.

## Declaration of interest

The authors declare no competing interests.

The full affiliations of Shingo Iwami are as follows: interdisciplinary Biology Laboratory (iBLab), Division of Natural Science, Graduate School of Science, Nagoya University, Nagoya, Japan; Institute of Mathematics for Industry, Kyushu University, Fukuoka, Japan; Institute for the Advanced Study of Human Biology (ASHBi), Kyoto University, Kyoto, Japan; Interdisciplinary Theoretical and Mathematical Sciences Program (iTHEMS), RIKEN, Saitama, Japan; NEXT-Ganken Program, Japanese Foundation for Cancer Research (JFCR), Tokyo, Japan; International Research Center for Neurointelligence, The University of Tokyo Institutes for Advanced Study, The University of Tokyo, Tokyo, Japan; Science Groove Inc., Fukuoka, Japan.

The full affiliations of Shoya Iwanami are as follows: interdisciplinary Biology Laboratory (iBLab), Division of Natural Science, Graduate School of Science, Nagoya University, Nagoya, Japan; Center for Surveillance, Immunization, and Epidemiologic Research, National Institute of Infectious Diseases, Tokyo, Japan

## Declaration of generative AI and AI-assisted technologies in the writing process

During the preparation of this work the authors used ChatGPT in order to improve the language of the manuscript. After using this tool, the authors reviewed and edited the content as needed and take full responsibility for the content of the publication.

## Notes

### Competing Interest Statement

The authors have declared no competing interest.

