## Supplementary Figures and Tables for "Mating status-dependent dopaminergic modulation of auditory sensory neurons in *Drosophila*"

Authors/affiliations

Haruna Yamakoshi[1], Mihoko Horigome[1], Shotaro Yamamoto[1], Shoya Iwanami[1], Shingo Iwami[1], Ryoya Tanaka[1], Yuki Ishikawa[1], & Azusa Kamikouchi[1, 2*]

1 Graduate School of Science, Nagoya University, Nagoya, Aichi, 464-8602, Japan

2 Institute of Transformative Bio-Molecules (WPI-ITbM), Nagoya University, Nagoya, Aichi, 464-8602, Japan

Author list footnotes

*Corresponding author

Contact info

This PDF file includes:

Supplementary Figures 1 to 17

Supplementary Tables 1 to 3

### Supplementary Figures

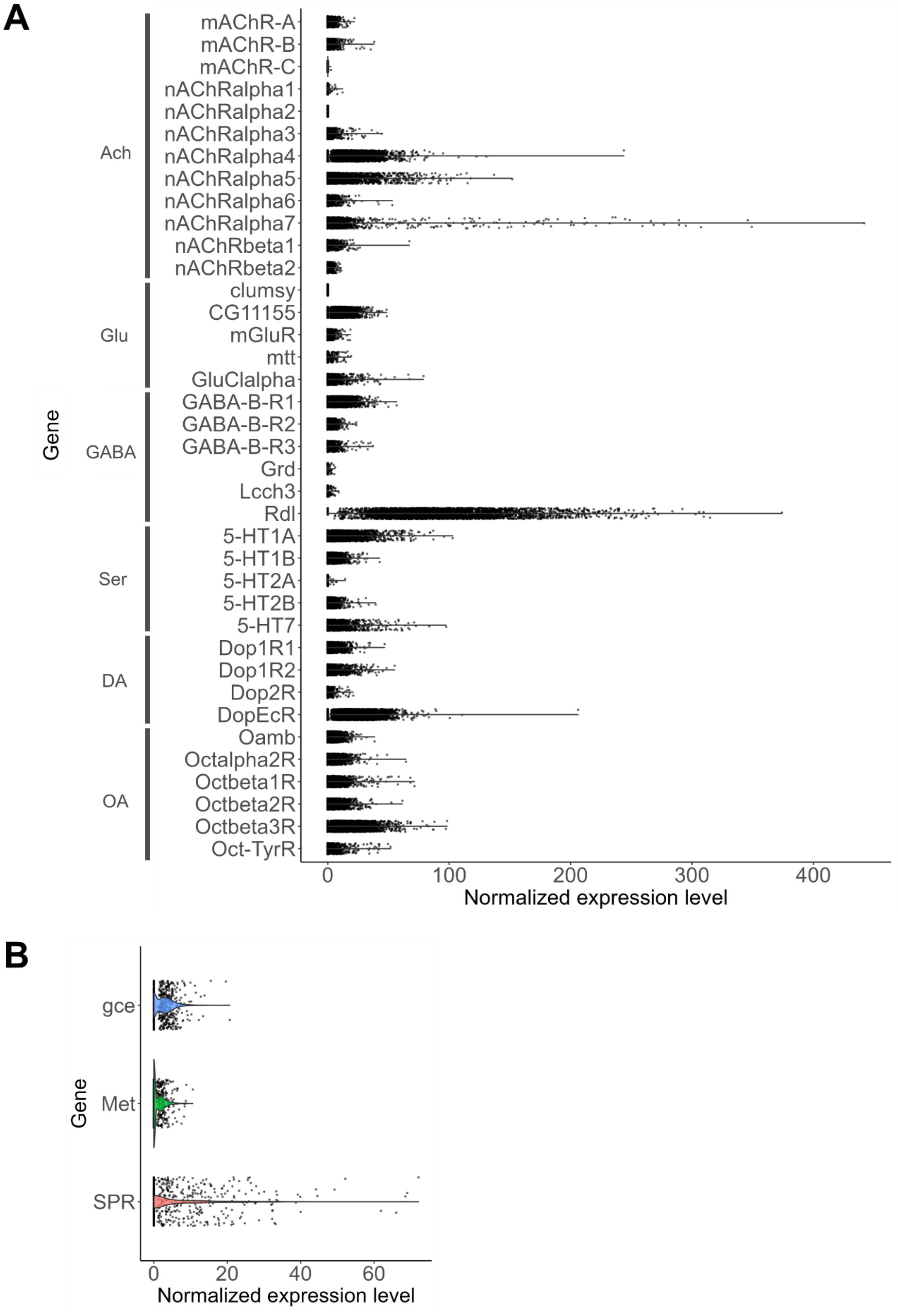

**Supplementary Figure S1: Normalized expression level of receptors in JO neurons.**

(A) Expression levels of neurotransmitter receptor gene in JO neurons. Data were obtained from the cell cluster annotated as JO neurons in the single-nucleus RNA-seq data.[1] *mAchR-A*, *mAchR-B*, *mAchR-C, nAchRalpha1, nAchRalpha2, nAchRalpha3, nAchRalpha4, nAchRalpha5, nAchRalpha6, nAchRalpha7, nAchRbeta1,* and *nAchRbeta2* are acetylcholine (Ach) receptors. *clumsy, CG11155*, *mGluR, mtt,* and *GluClalpha* are glutamate (Glu) receptors. *GABA-B-R1, GABA-B-R2*, *GABA-B-R3, Grd, Lcch3,* and *Rdl* are GABA receptors. *5-HT1A, 5-HT1B*, *5-HT2A, 5-HT2B,* and *5-HT7* are serotonin (Ser) receptors. *Dop1R1, Dop1R2*, *Dop2R,* and *DopEcR* are dopamine (DA) receptors. *Oamb, Octalpha2R, Octbeta1R, Octbeta2R, Octbeta3R,* and *Oct-TyrR* are octopamine (OA) receptors. No data was available for *nAchRbeta3, Nmdar1, Nmdar2, GluRIIA, GluRIIB, GluRIIC, GluRIID, GluRIIE, Ekar, GluRIA, GluRIB, Grik* and *KaiR1D.* JO, Johnston's organ.

(B) Expression levels of insect hormone receptor genes in *Dop1R2*-expressing JO neurons. Data were obtained from gene expression data of JO neurons in the single-nucleus RNA-seq data.[1] Because *Dop1R2* is expressed selectively in JO-AB neurons, we assume *Dop1R2*-expressing neurons from the JO neurons dataset are JO-AB neurons. *SPR*, sex peptide receptor. *gce* and *Met* encode juvenile hormone receptors.[2]

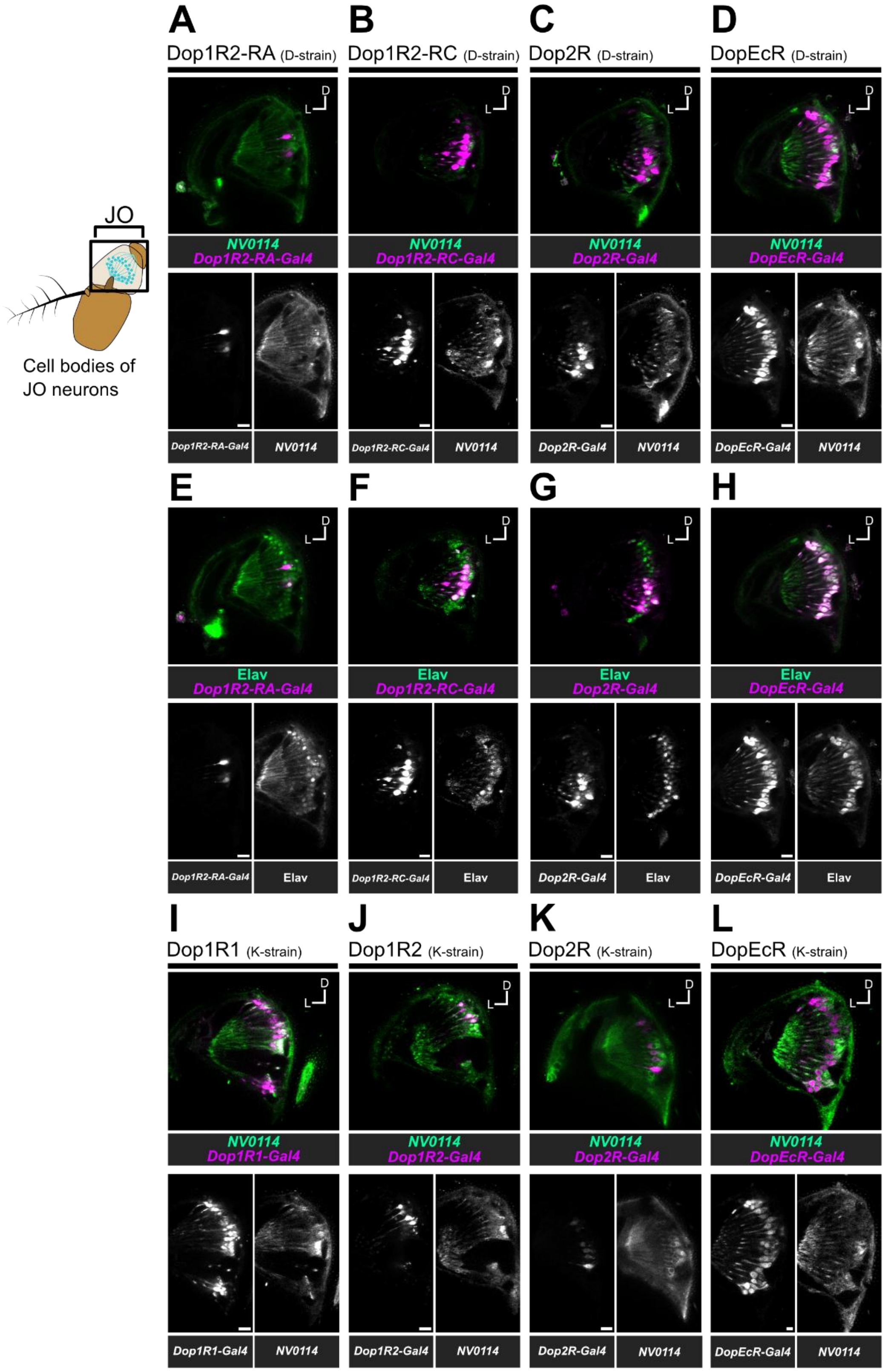


**Supplementary Figure S2: Expression patterns of T2A-Gal4 knock-in strains in JO.**

Expression patterns of dopamine receptors in JO. T2A-Gal4 knock-in strains reported by Deng et al.[3]

(A-H) and Kondo et al.[4] (I-L) are referred to as D-strain and K-strain, respectively. Labeling patterns of *Dop1R2-RA-Gal4* (A, E), *Dop1R2-RC-Gal4* (B, F), *Dop2R-Gal4* (C, G) and *DopEcR-Gal4* (D, H) from D-strain and *Dop1R1-Gal4* (I), *Dop1R2-Gal4* (J), *Dop2R-Gal4* (K) and *DopEcR-Gal4* (L) from K-strain are shown. T2A-Gal4 knock-in strains (magenta) were counter-labeled with *NV0114* LexA strain, which labels most JO neurons[5] (A-D, I-L) or Elav (a marker of neuronal cell bodies) (E-H) (green) in JO. Lower panels show the grayscale images of the top panels. The left diagrams show the position of JO. D, dorsal; L, lateral (the same in the following figures). Scale bar = 10 μm.

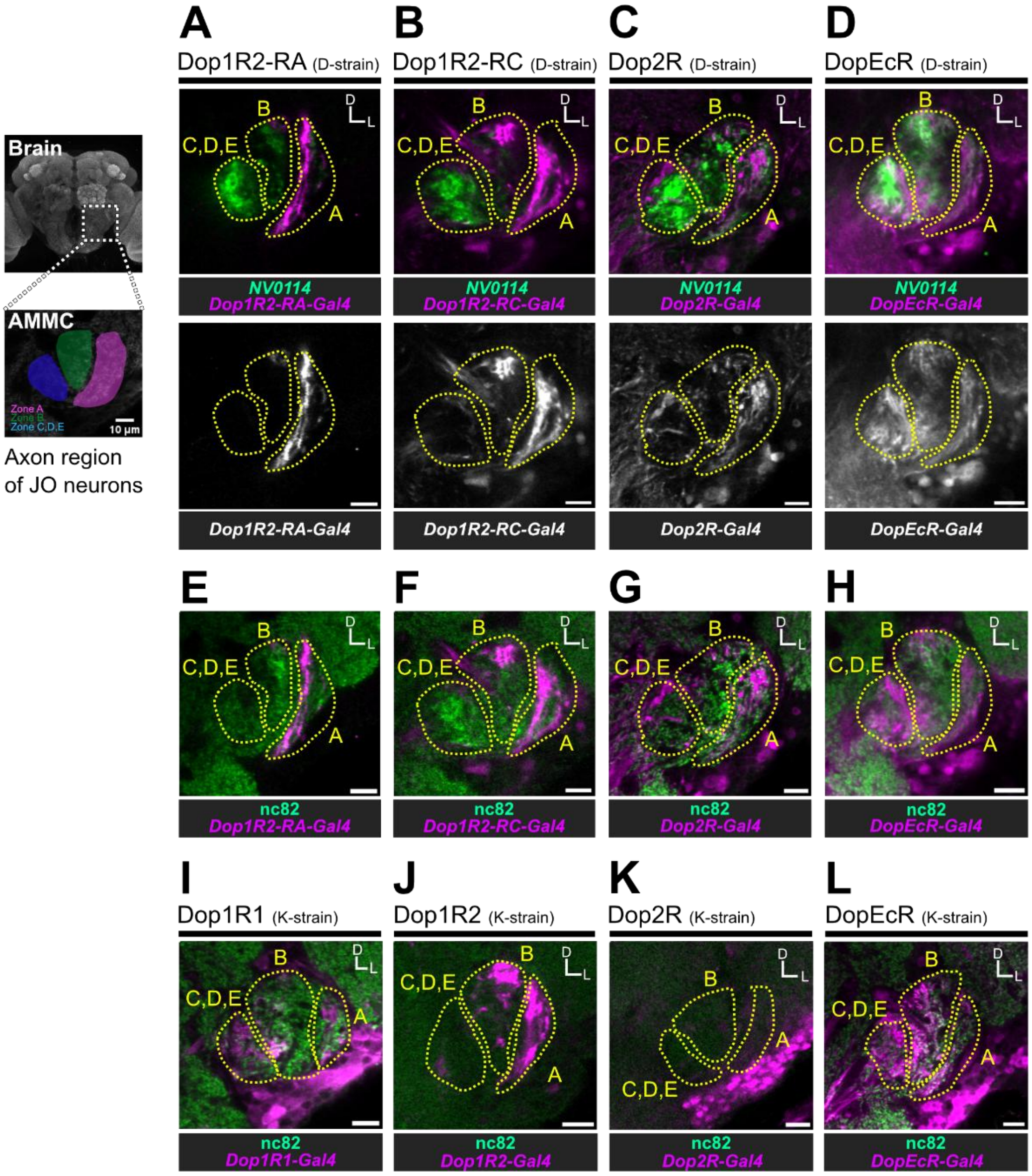


**Supplementary Figure S3: Expression patterns of T2A-Gal4 knock-in strains in the brain.**

Expression patterns of dopamine receptors in the brain. Labeling patterns of *Dop1R2-RA-Gal4* (A, E), *Dop1R2-RC-Gal4* (B, F), *Dop2R-Gal4* (C, G) and *DopEcR-Gal4* (D, H) from D-strains and *Dop1R1-Gal4* (I), *Dop1R -Gal4* (J), *Dop2R-Gal4* (K) and *DopEcR-Gal4* (L) from K-strains are shown. T2A-Gal4 knock-in strains (magenta) were counter-labeled with *NV0114* LexA strain, which labels most JO neurons[5] (A-D) or nc82 (synaptic marker) (E-L) (green) in JO. Lower panels show the grayscale images of the top panels (A-D). The left diagrams show the AMMC in the brain with each axon region of JO neurons, i.e., zones A to E. D, dorsal; L, lateral (the same in the following figures). AMMC, antennal mechanosensory and motor center. Scale bar = 10 µm.

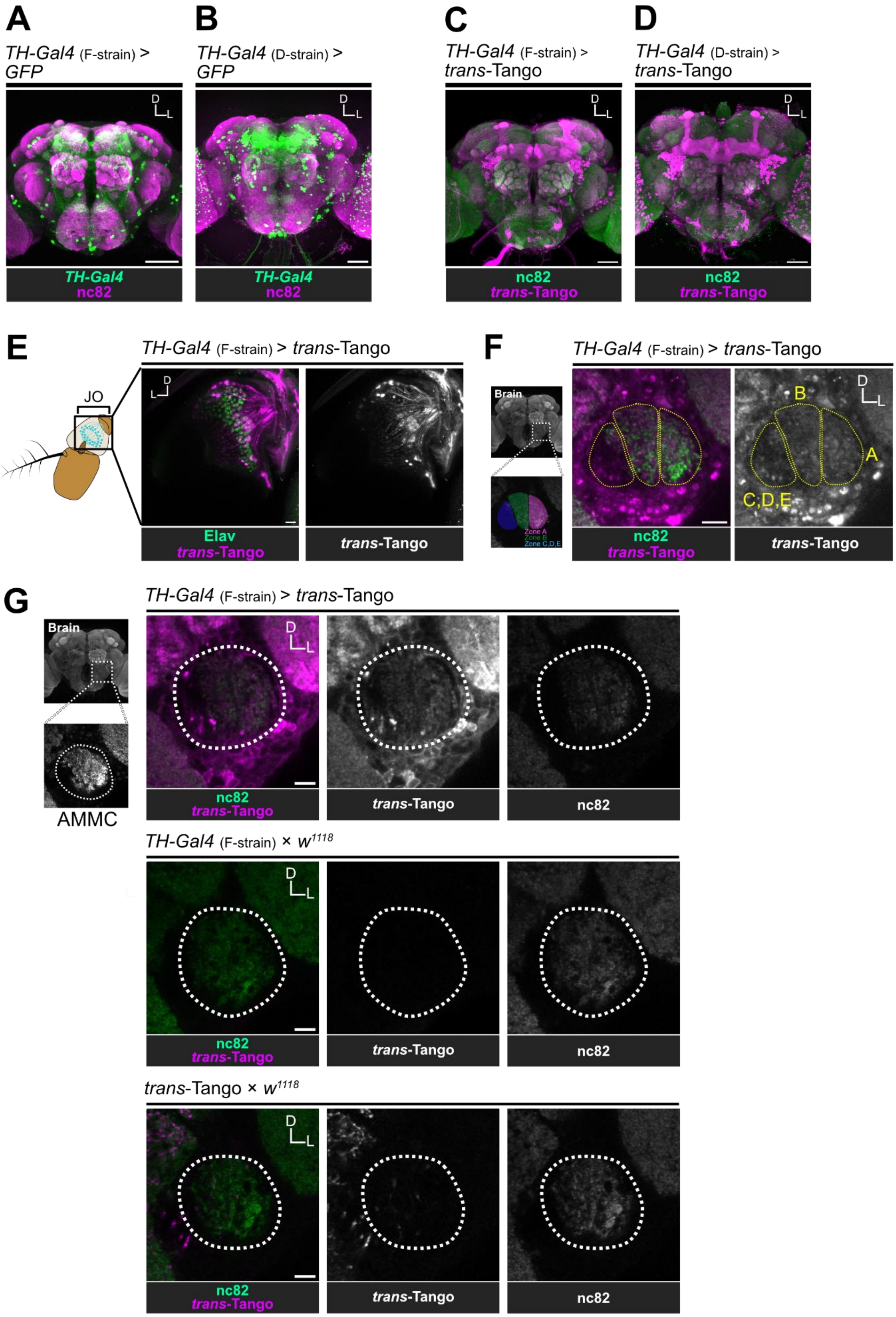
A
TH-Gal4 (F-strain) > GFP
TH-Gal4
nc82
B
TH-Gal4 (D-strain) > GFP
TH-Gal4
nc82
C
TH-Gal4 (F-strain) > trans-Tango
nc82
trans-Tango
D
TH-Gal4 (D-strain) > trans-Tango
nc82
trans-Tango
E
JO
TH-Gal4 (F-strain) > trans-Tango
Elav
trans-Tango
trans-Tango
F
Brain
Zone A
Zone B
Zone C,D,E
TH-Gal4 (F-strain) > trans-Tango
B
A
C,D,E
nc82
trans-Tango
trans-Tango
G
Brain
AMMC
TH-Gal4 (F-strain) > trans-Tango
nc82
trans-Tango
trans-Tango
nc82
TH-Gal4 (F-strain) × w1118
nc82
trans-Tango
trans-Tango
nc82
trans-Tango × w1118
nc82
trans-Tango
trans-Tango
nc82

67

**Supplementary Figure S4: Synaptic connections between *TH-Gal4* neurons and JO neurons**.

(A, B) Expression patterns of two *TH-Gal4* driver strains in the brain. Strains reported by Friggi-Grelin et al[6] (A) and Deng et al.[3] (B) are referred to as F-strain and D-strain, respectively. Labeled neurons in each strain are visualized by expressing mCD8::GFP markers (green). nc82 antibodies visualize neuropils (magenta). Scale bar = 50 µm.

(C, D) *trans*-Tango signals of *TH-Gal4* strains by Friggi-Grelin et al[6] (C) and Deng et al. [3] (D) in the central brain. tdTomato was used to visualize *trans*-Tango signals, labeled with anti-DsRed antibodies (magenta). nc82 antibodies visualize neuropils (green). Scale bar = 50 µm.

(E, F) *trans*-Tango signals of *TH-Gal4* strains in JO (C) and AMMC (D). *TH-Gal4* strain generated in Friggi-Grelin et al.[6] (F-strain) was used. Scale bar = 10 µm.

(G) *trans*-Tango signals in the AMMC of *TH-Gal4*>*trans*-Tango flies (top) and parental control flies (middle and bottom). *TH-Gal4* strain generated in Friggi-Grelin et al.[6] (F-strain) was used. *TH-Gal4*/ $w^{1118}$ flies (middle) and *trans*-Tango/ $w^{1118}$ flies (bottom) were used as control flies. tdTomato was used to visualize *trans*-Tango signals, labeled with anti-DsRed antibodies (magenta). nc82 antibodies visualize neuropils (green). Dotted white lines show the outline of the AMMC. Scale bar = 10 µm.

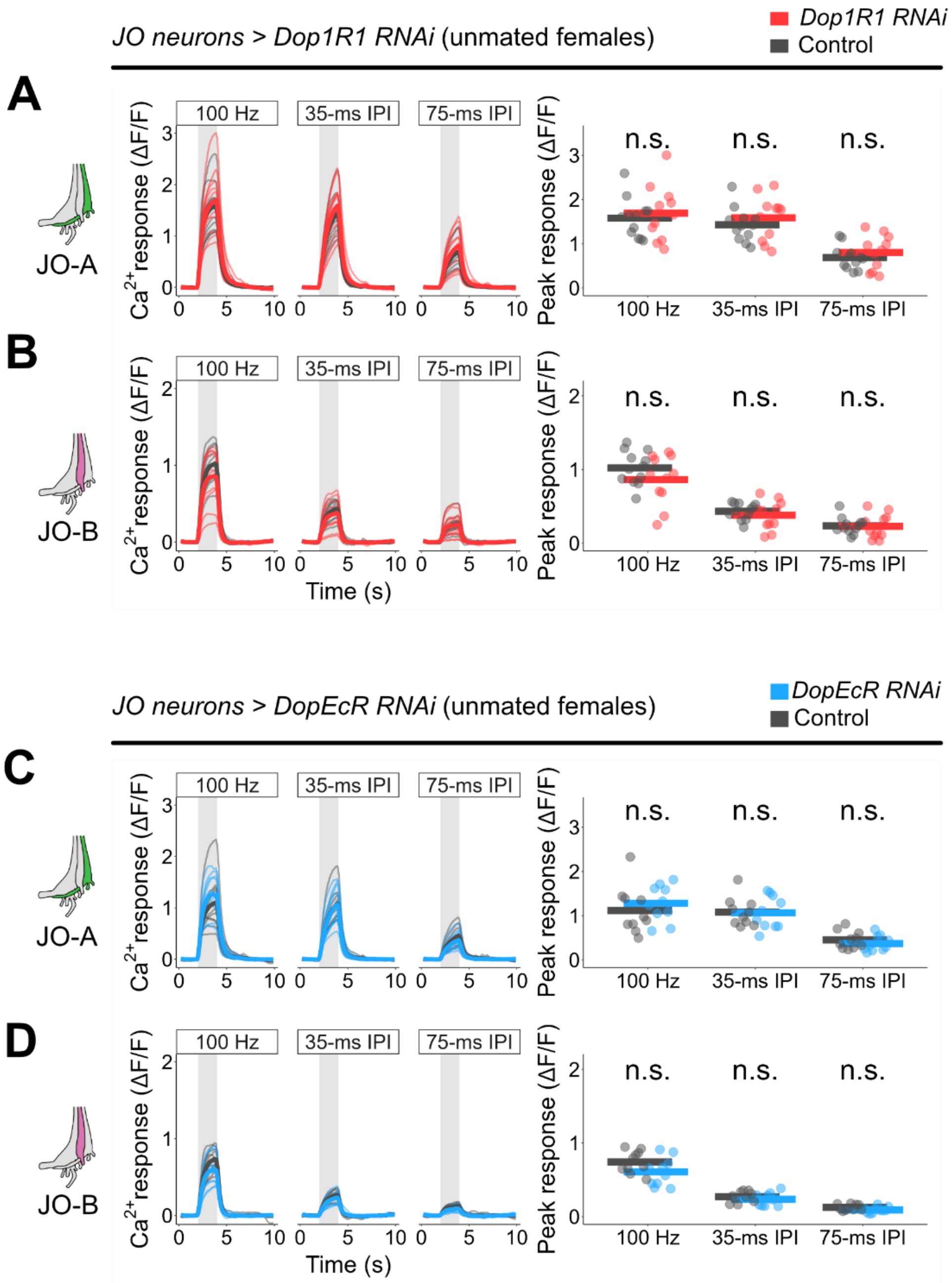


**Supplementary Figure S5: *Dop1R1* or *DopEcR* knockdown in JO neurons have no effect on calcium responses.**

Calcium responses were measured from unmated females with *Dop1R1* (A, B) or *DopEcR* (C, D) knockdown in JO neurons. Responses of JO-A neurons (A, C) and JO-B neurons (B, D) to a pure tone of 100 Hz and artificial pulse songs (35 ms and 75 ms IPI) are shown. Left, Time traces of raw ΔF/F responses. Gray-shaded areas indicate the time window of sound playback. Thin and bold lines show the responses in each individual and the average of all individuals, respectively Right, peak calcium responses. Dots and bars show the peak responses in each individual and the average of all individuals, respectively (the same in following figures). n = 10–12 per genotype.

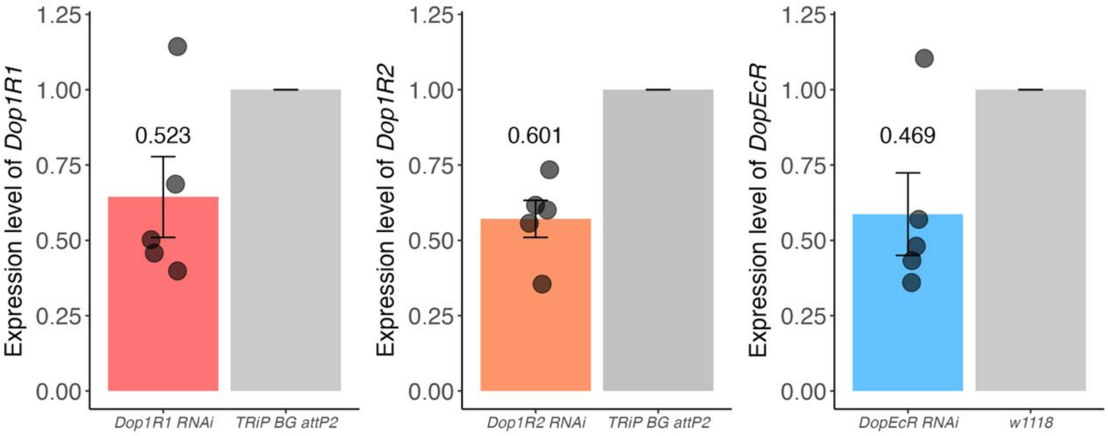


**Supplementary Figure S6: RNAi efficiency evaluated by RT-qPCR**

RNAi efficiency was assessed by quantitative RT-qPCR. Expression of each RNAi construct was driven by the *actin-Gal4* driver line. Transcript levels of RNAi knockdown groups were normalized to those in the control group (*TRiP BG attP2* or $w^{1118}$). Ribosomal protein 49 (*RP49*) was used as an internal control. Average relative expression levels are shown above the bar graphs. Error bars represent the standard error of the mean (SEM). The median relative expression for each graph is indicated above the bar graph. Each dot represents the relative expression level of each repeat. n = 5 for each graph.

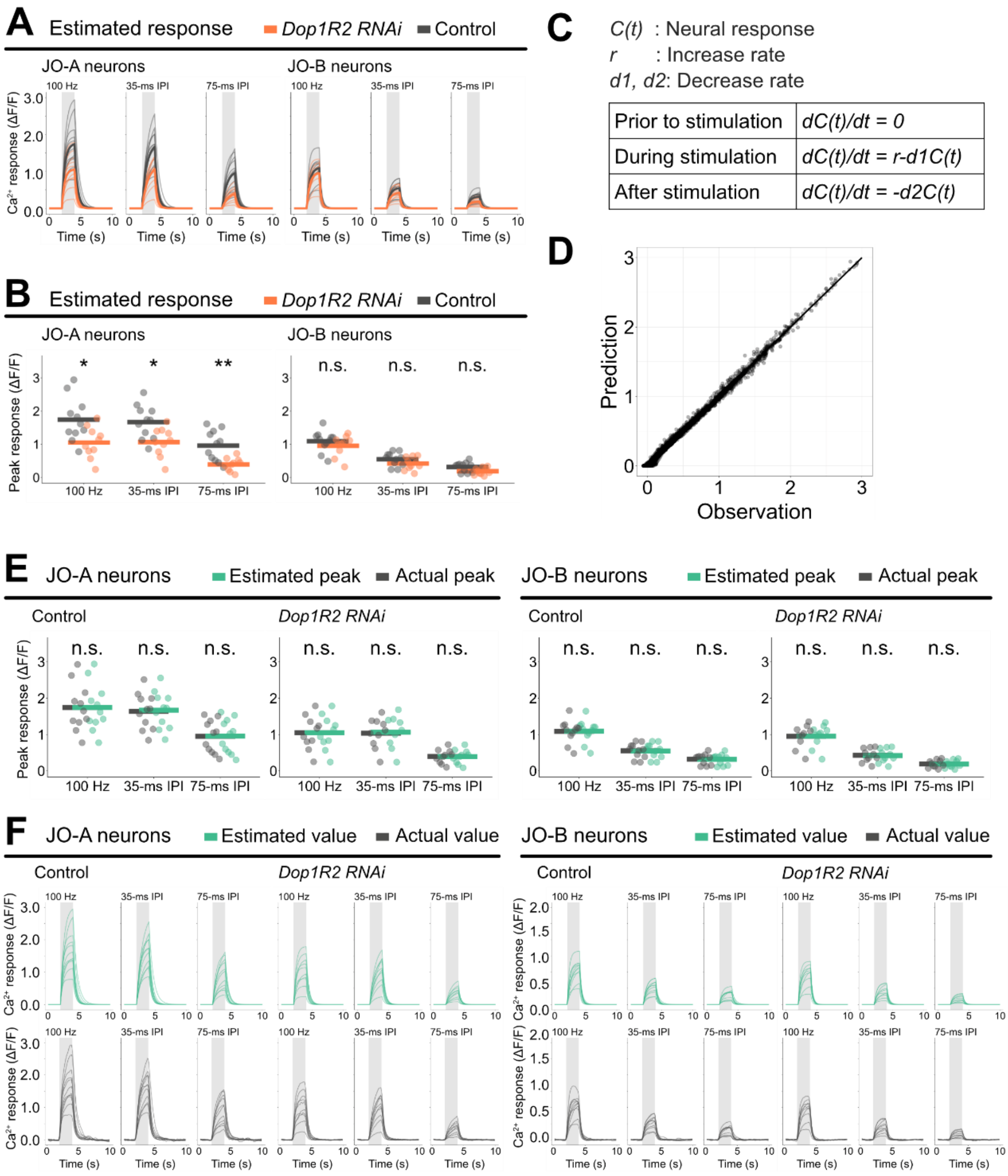


**Supplementary Figure S7: Validation of the three-component model that describes calcium dynamics**.

To elucidate which components of the calcium dynamics were affected by *Dop1R2* knockdown in JO-A neurons, we fitted the time series data of calcium responses to a simple three-component model that incorporates the following factors: the increase rate during the stimulus ($r$), and the decrease rates during ($d_1$) and after ($d_2$) the stimulus. The estimated calcium responses using this model well replicated the actual data, validating the model (Supplementary Table S2).

(A) Time traces of estimated calcium responses in JO-A and JO-B neurons. Estimations in *Dop1R2* knockdown (orange) and its control group (grey) are shown.

(B) Estimated peak calcium responses. Dots and bars show peak responses in each individual and the average of all individuals, respectively.

(C) Model used to estimate calcium responses. The increase rate ($r$) and two types of decrease rate ($d_1$, $d_2$) were used as the components of the model.

(D) Observed responses plotted against estimated responses. Each dot represents calcium response of one frame (0.1 s).

(E) Comparison between estimated (green) and actual peaks (grey).

(F) Time traces of estimated (green) and actual (grey) calcium responses.

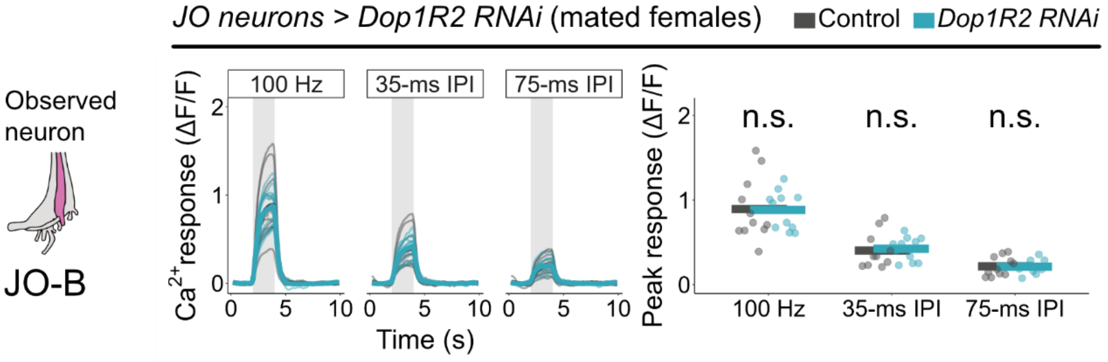


**Supplementary Figure S8: *Dop1R2* knockdown does not affect JO-B responses in mated females.**

Calcium responses in JO-B neurons to three types of sound stimuli were measured in mated females. n =11 per genotype.

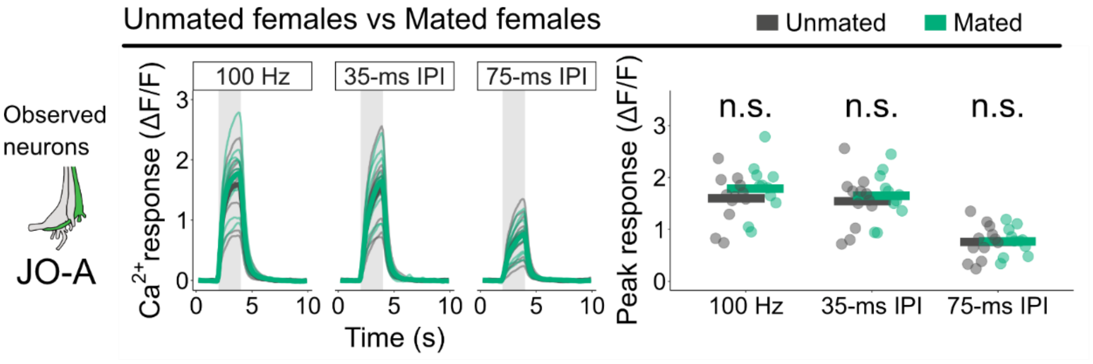


**Supplementary Figure S9: Calcium responses are similar across different mating states.**

Calcium responses in JO-A neurons to three types of sound stimuli were measured in unmated and mated females. No significant difference was detected in each comparison. n =11-12 per genotype.

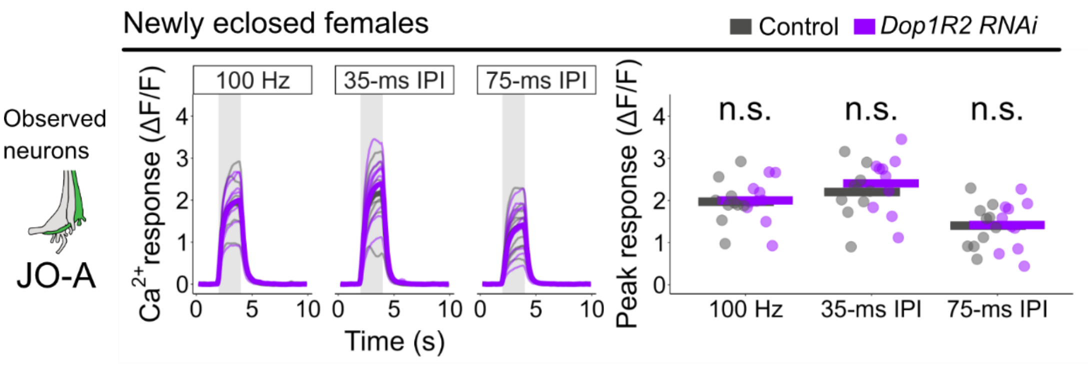


**Supplementary Figure S10: *Dop1R2* knockdown does not significantly affect the calcium responses of JO-A neurons in newly eclosed females.**

Calcium responses of JO-A neurons in newly eclosed females (26-36 hours after eclosion) with *Dop1R2* knockdown in JO neurons. n =10 per genotype.

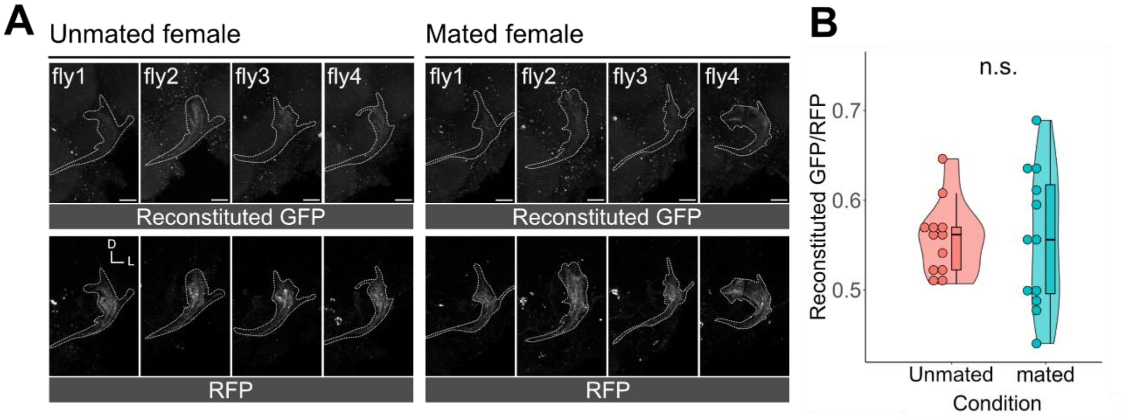


**Supplementary Figure S11: Similar expression levels of Dop1R2 between JO-A axons of unmated and mated females.**

(A) Reconstituted GFP (Top) and RFP signals (Bottom) in the brains of unmated (Left) and mated females (Right). The expressions of RFP and GFP$^{1\text{-}10}$ were driven by *R74C10* -Gal4, which selectively labels JO-A neurons (Figure S14). The endogenous Dop1R2 was tagged with GFP$^{11}$ at its C terminus. The brain areas around the AMMC of representative flies are shown. The regions expressing RFP signals are outlined with white dotted lines. Each image was z-stacked by max pixel intensity. Scale bar = 20 μm.

(B) The reconstituted GFP / RFP ratio indicating normalized GFP intensity. Average reconstituted GFP fluorescence normalized to RFP fluorescence along the JO-A neuron axons in each individual is represented as a dot. In the boxplot, a horizontal line within the box represents the median, while the vertical lines extending outside the box indicate the interquartile range. Violin plot illustrates the data distribution. n =12 per condition.

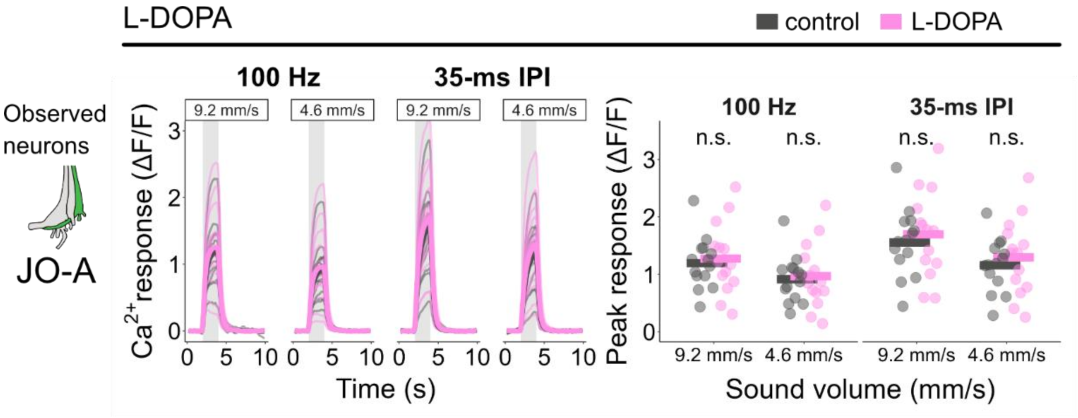


**Supplementary Figure S12: Manipulation of dopamine level does not affect the calcium responses to sound in mated females.**

Calcium responses in JO-A neurons to two types of sound stimuli were measured in L-DOPA-fed mated females. Responses were recorded at two different sound intensities (9.2 and 4.6 mm/s particle velocity, peak-to-peak amplitude). n =11-12 per genotype.

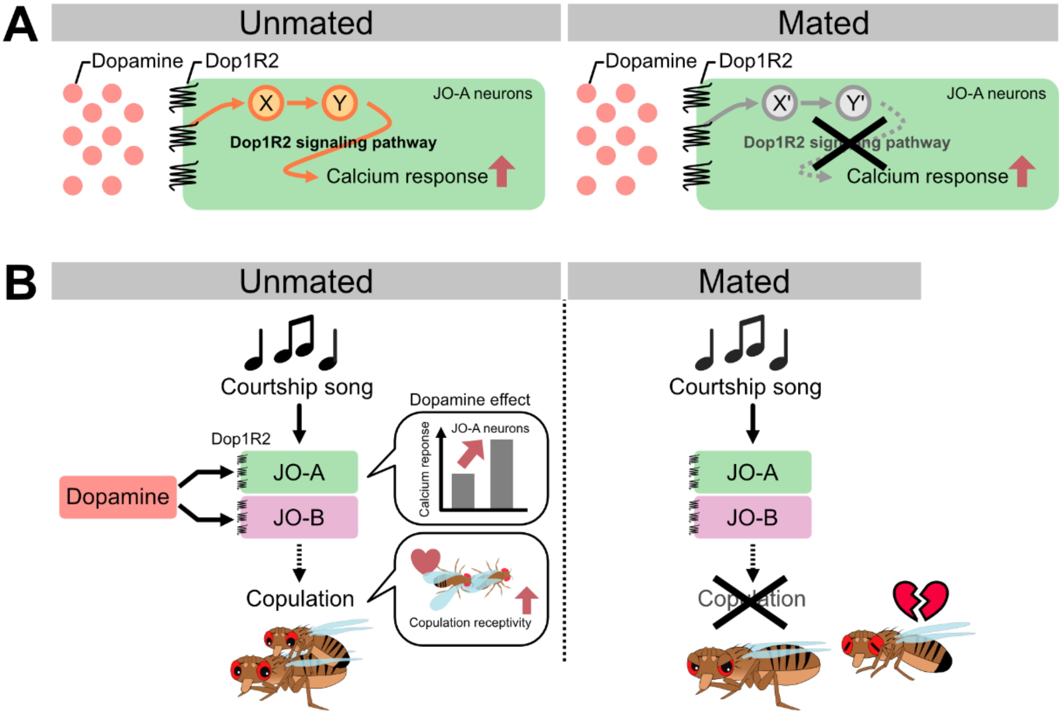


**Supplementary Figure S13: Possible models for mating-status dependent dopaminergic modulation of auditory sensory neurons in *Drosophila* females.**

(A) A model for mating-status dependent modulation of JO-A neurons. This study suggests that Dop1R2 expression level and upstream dopamine release are not to primary factors causing the difference between unmated and mated females (Figures S11, S12). In mated females, therefore, the intracellular signaling pathway downstream of Dop1R2 may be suppressed.

(B) A model for mating-status dependent modulation of auditory sensory neurons. Courtship songs emitted by males activate JO-A and JO-B neurons of females. Modulation via Dop1R2 enhances the calcium responses to sound in JO-A neurons. Moreover, expression of Dop1R2 in JO-A and JO-B neurons enhances female copulation receptivity when exposed to the song.

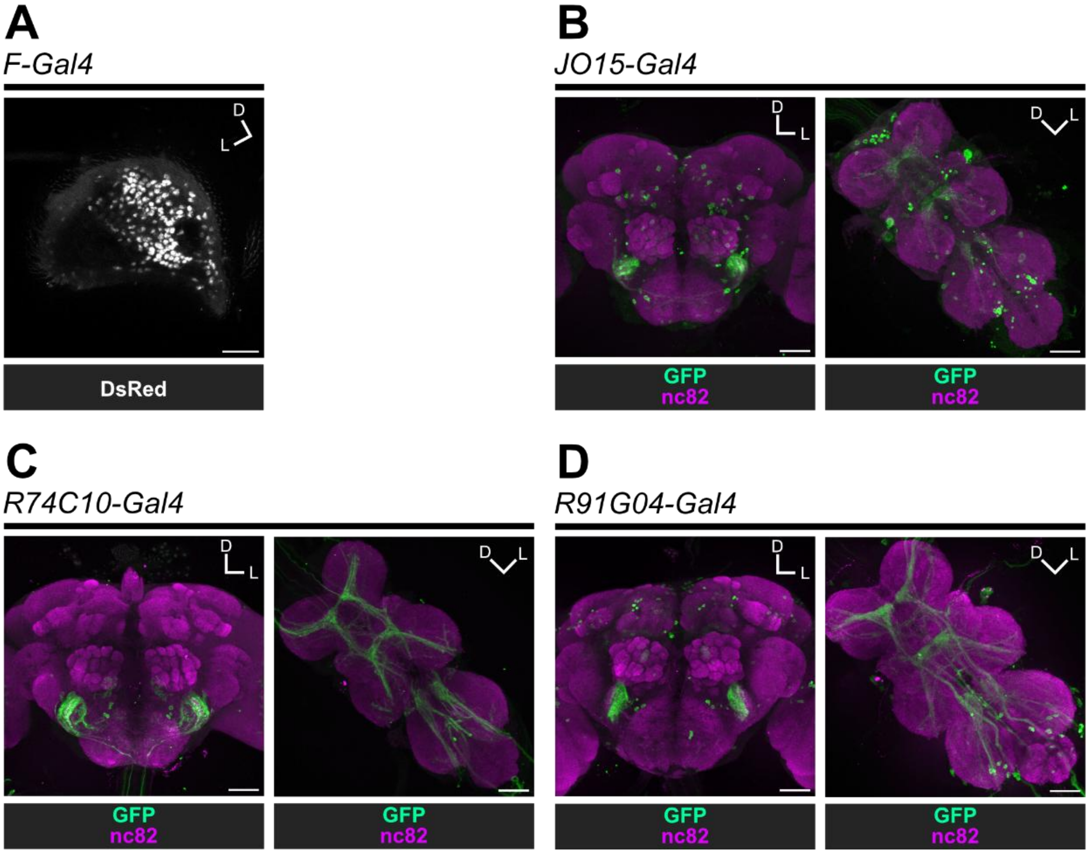


**Supplementary Figure S14: Expression patterns of GAL4 drivers used in the behavioral experiments.**

(A) Labeling patterns of *F-Gal4* in JO. The DsRed marker is expressed to visualize *F-Gal4* positive cells (white) in the second antennal segment where the JO resides. These cell bodies of JO neurons send their axons to the brain.[7] Scale bar = 20 $\mu$m.

(B-D) Labeling patterns in the brain (Left) and ventral nerve cord (Right). *JO15-Gal4* (B), *R74C10-Gal4* (C), and *R91G04-Gal4* (D) were used to express mCD8::GFP markers (green). nc82 antibodies were used for counter-labeling (magenta). Scale bar = 50 $\mu$m.

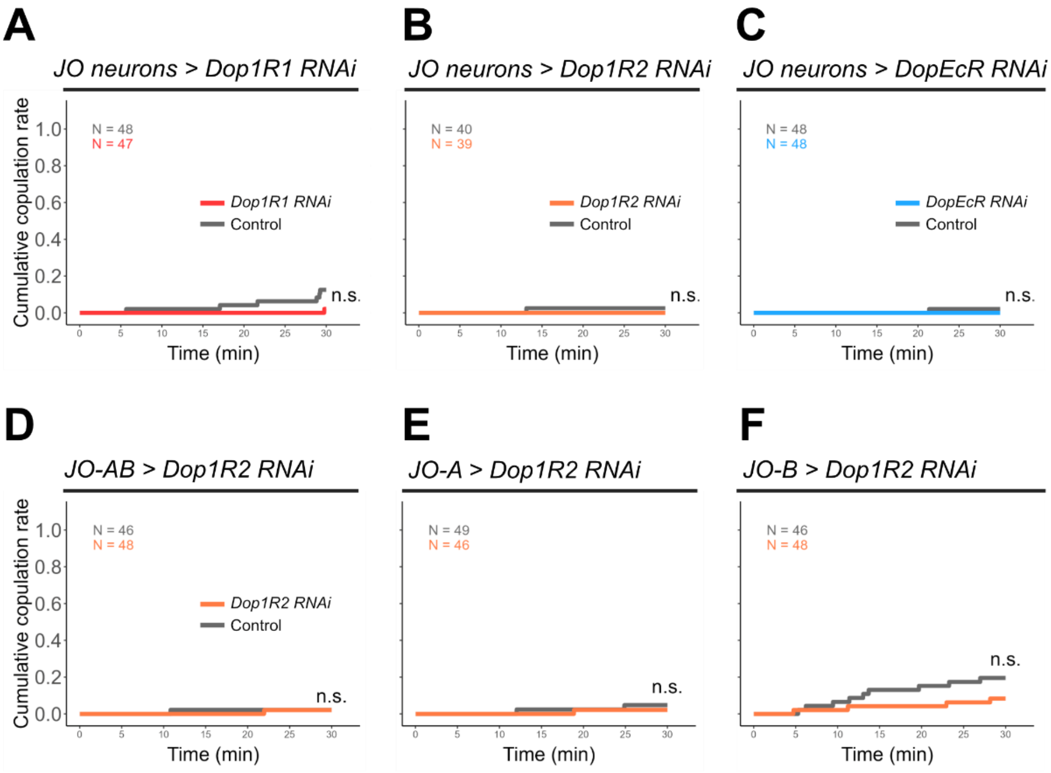


**Supplementary Figure S15: Knockdown of dopamine receptors does not affect female copulation receptivity without song exposure.**

Cumulative copulation rates in no sound conditions are shown in each panel.

(A-C) Females with *Dop1R1* (A), *Dop1R2* (B), or *DopEcR* (C) knockdown in JO neurons were tested. *F-Gal4* driver strain was used to express each RNAi construct.

(D-F) Females with *Dop1R2* knockdown in JO-AB neurons (D), JO-A neurons (E) and JO-B neurons (F) were tested. Driver strains used in each panel are the same as those used in Figures 5, 6, and S16.

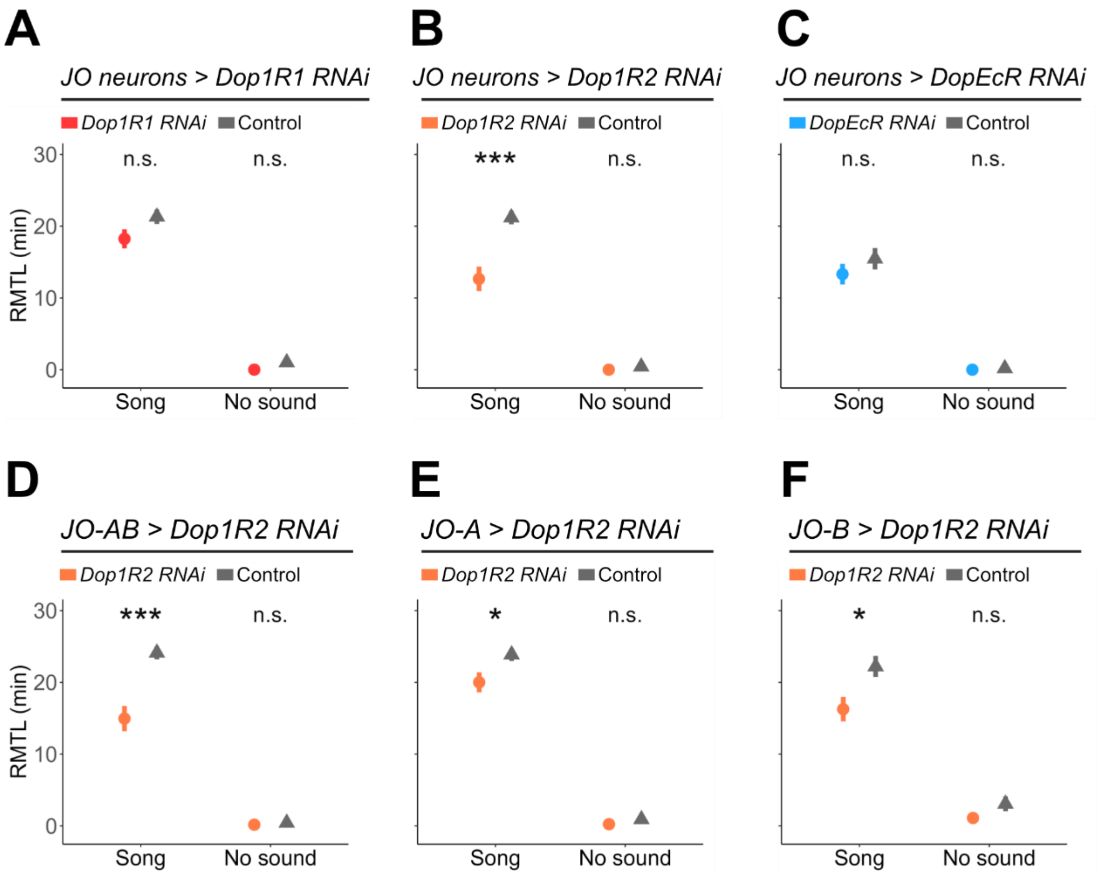


**Supplementary Figure S16: Evaluation of female receptivity by restricted mean time lost (RMTL).**

RMTLs of females with gene knockdowns and their controls are shown.

(A-C) *Dop1R1* (A), *Dop1R2* (B) and *DopEcR* (C) were knocked down in JO neurons.

(D-F) *Dop1R2* were knocked down in JO-AB neurons (D), JO-A neurons (E) and JO-B neurons (F).

Dot and vertical line represent the estimated value and standard error, respectively.

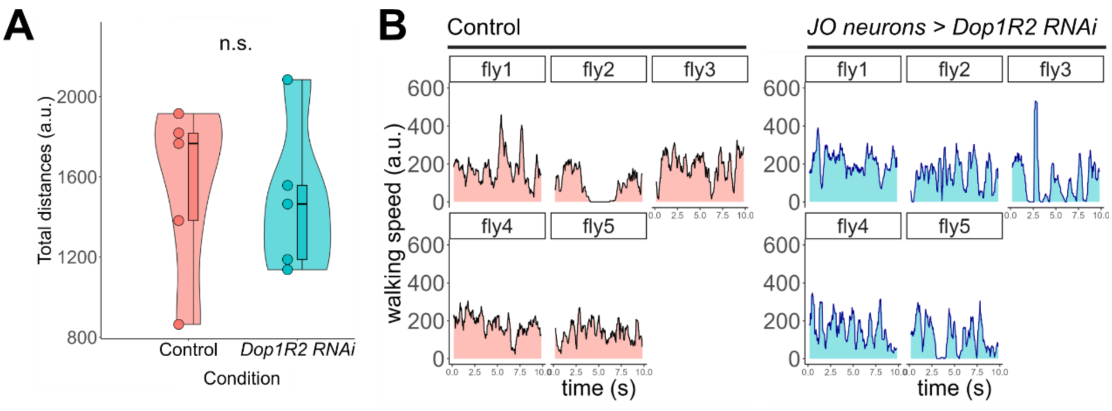


**Supplementary Figure S17: Locomotor activity of females during copulation assays.**

The total walking distances (A) and speeds (B) within 10 sec (300 frames) following the onset of male chasing behavior in control (left) and *F-Gal4>Dop1R2 RNAi* (right) females. The analysis used the five pairs with the earliest mating initiation times in each group.

**Table S1: Fly genotype.**

**Males**

| Figure | Genotype |
|---|---|
| 4B, 5, 6, S8, S9, S11, S12, S15, S16, S17 | *Canton-S* |
| 4C | *SP0/ Df(3L)Δ130* |

**Females**

| Figure | Genotype |
|---|---|
| 1B, 1F, S2I, S3I | *NV0116; UAS-DsRed (C5), lexAop-rCD2::GFP/ Dop1R1-Gal4*[4] |
| 1C, 1G, S2J, S3J | *NV0116; UAS-DsRed (C5), lexAop-rCD2::GFP/ Dop1R2-Gal4*[4] |
| S2A, S2E, S3A, S3E | *NV0116; UAS-DsRed (C5), lexAop-rCD2::GFP/ Dop1R2-RA-Gal4*[3] |
| S2B, S2F, S3B, S3F | *NV0116; UAS-DsRed (C5), lexAop-rCD2::GFP/ Dop1R2-RC-Gal4*[3] |
| 1D, 1H, S2K, S3K | *Dop2R-Gal4*[4]; *NV0116; UAS-DsRed (C5), lexAop-rCD2::GFP/ +* |
| S2C, S2G, S3C, S3G | *Dop2R-Gal4*[3] ; *NV0116; UAS-DsRed (C5), lexAop-rCD2::GFP/+* |
| 1E, 1I, S2L, S3L | *NV0116; UAS-DsRed (C5), lexAop-rCD2::GFP/DopEcR-Gal4*[4] |
| S2D, S2H, S3D, S3H | *NV0116; UAS-DsRed (C5), lexAop-rCD2::GFP/DopEcR-Gal4*[3] |
| 2B, 2C, S4B, S4D | *trans-Tango, QUAS-mtdTomato-3xHA; UAS-IVS-mCD8::GFP/TH-Gal4*[3] |
| S4A, S4C, S4E, S4F, S4G (top) | *trans-Tango, QUAS-mtdTomato-3xHA; UAS-IVS-mCD8::GFP/TH-Gal4*[6] |
| S4G (middle) | *w1118;;TH-Gal4*[6] |
| S4G (bottom) | *w1118; trans-Tango, QUAS-mtdTomato-3xHA; UAS-IVS-mCD8::GFP/+* |
| 3, 4, S7, S8, S10 | *F-Gal4, UAS-GCaMP6f; iav-Gal4/ Dop1R2 RNAi* |
| | *F-Gal4, UAS-GCaMP6f; iav-Gal4/ TRiP* background line (Control) |
| S5A, S5B | *F-Gal4, UAS-GCaMP6f; iav-Gal4/ Dop1R1 RNAi* |
| | *F-Gal4, UAS-GCaMP6f; iav-Gal4/ TRiP* background line (Control) |
| S5C, S5D | *F-Gal4, UAS-GCaMP6f/DopEcR-RNAi; iav-Gal4* |
| | *F-Gal4, UAS-GCaMP6f/w1118; iav-Gal4* (Control) |
| S6 (left) | *y[1] w[*]/y[1] v[1]; actin-Gal4; Dop1R1 RNAi* |
| | *y[1] w[*]/y[1] v[1]; actin -Gal4;TRiP* background line (Control) |
| S6 (center) | *y[1] w[*]/y[1] v[1]; actin -Gal4; Dop1R2 RNAi* |
| | *y[1] w[*]/y[1] v[1]; actin -Gal4; TRiP* background line (Control) |
| S6 (right) | *actin -Gal4/DopEcR RNAi;* |
| | *actin -Gal4/w1118* (Control) |
| S9 | *F-Gal4, UAS-GCaMP6f; iav-Gal4/ TRiP* background line |

| | |
|---|---|
| S11 | *UAS-GFP1-10/10XUAS-IVS-mCD8::RFP; Dop1R2-RA-7xGFP11/ R74C10-Gal4* |
| S12 | *F-Gal4, UAS-GCaMP6f/+ ; iav-Gal4/+* |
| Table 9 | *UAS-GFP1-10/10XUAS-IVS-mCD8::RFP;+ / R74C10-Gal4* |
| 5C, 5F, S15A, S16A | *w[*]/y[1] v[1]; F-Gal4; Dop1R1 RNAi* |
| | *w[*]/y[1] v[1]; F-Gal4;TRiP* background line (Control) |
| 5D, 5F, 6E, S15B, S16B, S17 | *w[*]/y[1] v[1]; F-Gal4; Dop1R2 RNAi* |
| | *w[*]/y[1] v[1]; F-Gal4; TRiP* background line (Control) |
| 5E, 5F, S15C, S16C | *F-Gal4/DopEcR RNAi;* |
| | *F-Gal4/w1118* (Control) |
| 6A, 6D, 6E, S15D, S16D | *w[*]/y[1] v[1];;JO15-Gal4/Dop1R2 RNAi* |
| | *w[*]/y[1] v[1];;JO15-Gal4/ TRiP* background line (Control) |
| 6B, 6D, 6E, S15E, S16E | *w[1118]/y[1] v[1];;R74C10-Gal4/Dop1R2 RNAi* |
| | *w[1118]/y[1] v[1];; R74C10-Gal4/ TRiP* background line (Control) |
| 6C, 6D, 6E, S15F, S16F | *w[1118]/y[1] v[1];;R91G04-Gal4/Dop1R2 RNAi* |
| | *w[1118]/y[1] v[1];; R91G04-Gal4/ TRiP BG attP2* (Control) |
| 5B | *w[*]; F-Gal4; 20XUAS IVS mCD8::GFP* |
| S14A | *F-Gal4/ NV0116; UAS-DsRed (C5), lexAop-rCD2::GFP/ +* |
| S14B | *w[*];; JO15-Gal4/ 20XUAS IVS mCD8::GFP* |
| S14C | *w[1118]/ w[*];;R74C10-Gal4/ 20XUAS IVS mCD8::GFP* |
| S14D | *w[1118]/ w[*];; R91G04-Gal4/ 20XUAS IVS mCD8::GFP* |

**Table S2: Statistical analysis of predicted calcium responses.**

| Figure | Strain | Female state | Sound | Neuron | t-value | p-value | Adjusted p-value | Cohen's d |
|---|---|---|---|---|---|---|---|---|
| S7B | *F-Gal4, iav-Gal4 >Dop1R2 RNAi, GCaMP6f* | Unmated | 100 Hz | JO-A | 2.81 | 0.0117 | 0.0117 | 1.21 |
| | | | 35-ms IPI | | 2.94 | 0.00835 | 0.0117 | 1.28 |
| | | | 75-ms IPI | | 3.72 | 0.00238 | 0.00714 | 1.57 |
| S7B | | | 100 Hz | JO-B | 1.01 | 0.324 | 0.324 | 0.442 |
| | | | 35-ms IPI | | 1.50 | 0.150 | 0.225 | 0.653 |
| | | | 75-ms IPI | | 2.30 | 0.00334 | 0.100 | 0.990 |
| S7E | *F-Gal4, iav-Gal4> TRiP* background, *GCaMP6f* | Unmated | 100 Hz | JO-A | -0.00246 | 0.998 | 0.998 | -0.00105 |
| | | | 35-ms IPI | | -0.149 | 0.883 | 0.998 | -0.0634 |
| | | | 75-ms IPI | | -0.0497 | 0.961 | 0.998 | -0.0212 |
| S7E | *F-Gal4, iav-Gal4* | Unmated | 100 Hz | JO-B | -0.0510 | 0.960 | 0.965 | -0.0218 |

| | | | | | | | | |
|---|---|---|---|---|---|---|---|---|
| | *>Dop1R2 RNAi, GCaMP6f* | | 35-ms IPI | | 0.0450 | 0.965 | 0.965 | 0.0192 |
| | | | 75-ms IPI | | 0.106 | 0.917 | 0.965 | 0.0450 |
| S7E | *F-Gal4, iav-Gal4> TRiP* background, *GCaMP6f* | Unmated | 100 Hz | JO-A | 0.00228 | 0.998 | 0.998 | 0.00102 |
| | | | 35-ms IPI | | -0.142 | 0.888 | 0.998 | -0.0637 |
| | | | 75-ms IPI | | -0.0462 | 0.964 | 0.998 | 0.0207 |
| S7E | *F-Gal4, iav-Gal4 >Dop1R2 RNAi, GCaMP6f* | Unmated | 100 Hz | JO-B | -0.0285 | 0.978 | 0.978 | -0.0127 |
| | | | 35-ms IPI | | 0.0523 | 0.959 | 0.978 | 0.0234 |
| | | | 75-ms IPI | | 0.0888 | 0.930 | 0.978 | 0.0397 |

**Table S3: Primer design for RT-qPCR.**

| Gene | Primer Direction | Sequence | Primer length (bp) | Amplicon length (bp) |
|---|---|---|---|---|
| *Dop1R1* | Forward | AGGATCCGCTCAGATACGGC | 20 | 95 |
| | Reverse | CACAAAGGAGACGAACGCGG | 20 | |
| *Dop1R2* | Forward | TGGTCAACCTGCTGTCTGGG | 20 | 149 |
| | Reverse | CACAAAGGCCCTGCGAAAGTC | 21 | |
| *DopEcR* | Forward | GCCAACTTCAAGGGACCCAC | 20 | 123 |
| | Reverse | CCATTCTCCGGTCAAAGCGG | 20 | |
| *RP49* | Forward | AGTATCTGATGCCCAACATCG | 21 | 180 |
| | Reverse | CAATCTCCTTGCGCTTCTTG | 20 | |
